# Lipid Acyl Chain-Driven α-Synuclein Fibril Polymorphisms and Neuronal Pathologies

**DOI:** 10.1101/2025.09.10.675439

**Authors:** Yoongyeong Baek, Anika Alim, Yanheng Dong, Tarek Olabi, Jungwook Paek, Myungwoon Lee

## Abstract

Conformational variations in α-syn fibrils are thought to underlie the distinct clinical features of synucleinopathies, including Lewy body dementia (LBD), Parkinsons’s disease (PD), and multiple system atrophy (MSA), suggesting that distinct fibril structures act as molecular fingerprints linked to disease phenotype. While the origins of these conformational variations remain unclear, increasing evidence points to membranes as key modulators of fibrils conformations. In this study, we investigated how age-related alterations in membrane composition and fluidity influence α-syn fibril formation and cellular outcomes. Using complex mixture membranes that mimic normal neuronal membranes and their age-related modifications in fatty acid chains, we found that α-syn fibrils grown with these membranes displayed distinct 2D ssNMR spectral patterns compared to lipid-free α-syn fibrils, reflecting differences in rigid fibril cores. Moreover, fibrils grown with age-related membranes exhibited weaker membrane association than those grown with normal neuronal membranes. These membrane-associated fibrils induce stronger neuronal pathologies than lipid-free fibrils, though the severity differed in intraneuronal aggregation and inflammation responses. Overall, our findings provide new insights into how age-related changes in membrane composition shape α-syn fibril structure and pathogenicity, strengthening the link between membrane dynamics and amyloid-driven neurodegeneration.

## Introduction

The aggregation of misfolded α-synuclein (α-syn), characterized by its transition from a native conformation to β-sheet-rich amyloid fibrils, constitutes the primary component of cytoplasmic inclusions known as Lewy bodies (LBs)^1,2^. These LBs are key pathological hallmarks of synucleinopathies, including Parkinson’s disease (PD), Lewy body dementias (LBD), and multiple system atrophy (MSA)^3–8^. While synucleinopathies are typically classified based on clinical symptoms, the distribution of α-syn aggregates, and the affected cell types, growing research suggests that distinct α-syn fibril conformations are closely associated with specific synucleinopathies and contribute to their heterogeneous pathological features^9–13^. Nevertheless, the precise molecular mechanisms underlying α-syn fibril polymorphisms and their impact on synucleinopathy variability remain largely unknown.

α-Syn fibril formation is significantly influenced by multiple factors, including α-syn concentration^14,15^, temperature^16–18^, and interactions with small molecules such as RNA, DNA, and lipids^19–26^. These factors have been shown to modulate the structural characteristics of α-syn fibrils. Among them, lipids are recognized as key regulators of α-syn aggregation due to their strong interactions with α-syn in both physiological and pathological contexts^22–26^. In its native state, α-syn exists as a monomer and plays a crucial role in neurotransmitter release, particularly through its involvement in synaptic vesicle trafficking and fusion^27–29^. During these processes, α-syn is proposed to interact dynamically with various cellular membranes, including the neuronal plasma membrane and synaptic vesicles^29,30^. However, pathological conditions may alter lipid-α-syn interactions, contributing to LB accumulation. Because LBs are composed largely of lipids together with α-syn aggregates^31^, lipid interactions may catalyze the aggregation process, disrupt membrane structure, and promote co-aggregation. Consequently, lipids are now increasingly recognized as significant factors in modulating α-syn aggregation and its associated neurotoxicity.

Numerous studies have demonstrated that lipids modulate both the kinetics and conformation of α-syn fibrils^25,32–38^. For example, α-syn amyloid fibril formation is accelerated in the presence of anionic phospholipids such as phosphatidylglycerol (PG), phosphatidylserine (PS), or phosphatidic acid (PA) at relatively low lipid-to-protein (L/P) ratios^36,38–40^. This enhancement is attributed to electrostatic interactions that increase α-syn binding to membrane surfaces, thereby elevating its local α-syn monomer concentration and facilitating fibril nucleation. In contrast, binding to zwitterionic lipid vesicles composed of phosphatidylcholine (PC) either inhibits fibril formation or has minimal effect across various L/P ratios^37,41,42^. Beyond modulating fibril formation kinetics, membrane also influences α-syn fibril structures, as revealed by atomic force microscopy infrared (AFM-IR) spectroscopy, solid-state NMR (ssNMR), and cryo-electron microscopy (cryo-EM)^43–48^. Collectively, these studies highlight the critical role of lipid membranes in modulating α-syn amyloid formation.

However, despite these insights, most studies have relied on simplified membrane models containing only one or two lipid types, which do not fully recapitulate the complexity of neuronal membranes. Neuronal plasma membranes are primarily composed of multiple classes of phospholipids, cholesterol, and sphingolipids^40,49^. These complex lipid environments are essential for maintaining membrane integrity and regulating cellular function through interactions with diverse biomolecules. However, aging disrupts the lipid balance of neuronal membranes—most notably through a decline in polyunsaturated fatty acids (PUFAs) and an increase in monounsaturated fatty acids^50–52^. Such age-related lipid alterations reduce membrane fluidity, which may, in turn, affect α-syn membrane binding affinity and aggregation behavior, potentially influencing pathological outcomes. Yet, the impact of fatty acid-driven changes in membrane properties on α-syn aggregation and its subsequent neurotoxicity remains largely unexplored. Addressing this gap requires comprehensive structural and functional studies of α-syn fibrils formed in the presence of physiologically relevant plasma membranes representing both normal and aged conditions.

In this study, we investigated how membrane composition, particularly age-related changes in fatty acid saturation, affects α-syn aggregation. To this end, we developed two model membranes representing the plasma membranes of normal and aged neurons, referred to as Neuron and Aged membranes respectively. Both membrane models were composed of PC, phosphatidylethanolamine (PE), cholesterol, and sphingomyelin (SM) at a simplified molar ratio of 35:20:35:10. This composition was based on reported lipid abundances in neuronal plasma membranes and lipid profiling from brain tissue^40,53^. For the Neuron membranes, we selected 1-palmitoyl-2-oleoyl-glycero-3-phosphocholine (POPC, 16:0/18:1) and 2-dioleoyl-sn-glycero-3-phosphoethanolamine (DOPE, 18:1) to reflect the predominant fatty acid chain lengths and degrees of unsaturation commonly found in neuronal membranes^51,53,54^. To model age-associated changes, the Aged membranes was designed with reduced lipid unsaturation: PC was modified from POPC (16:0.18:1) to 1,2-dipalmitoyl-sn-glycero-3-phosphocholine (DPPC,16:0), and PE was modified from DOPE (18:1) to 1-palmitoyl-2-oleoyl-sn-glycero-3-phosphoethanolamine (POPE, 16:0/18:1), reflecting reported decreases in lipid unsaturation during aging^50,55,56^.

We then characterized α-syn fibril formation in the presence of Neuron and Aged membranes. Our results demonstrate that both membranes facilitate fibril formation, with the extent of acceleration depending on the L/P ratios. Moreover, each membrane environment generates distinct fibril structures with different membrane associations. Importantly, these structural differences in α-syn fibrils drive distinct intraneuronal aggregation patterns, neurodegeneration, and inflammatory responses, providing insight into how membrane composition contributes to the conformation-dependent pathological effects of α-syn fibrils.

## Materials and Methods

### Protein Expression

Recombinant full-length α-syn was expressed in *Escherichia coli (E. coli)* BL21 (DE3) competent cells. A single colony was inoculated into 10 mL of Luria-Bertani (LB) medium containing ampicillin (100 µg/ml) and incubated overnight at 37 °C with shaking at 150 rpm. The next day, the 10 mL culture was transferred into 1 L of LB medium containing ampicillin (100 µg/ml) and incubated at 37 °C with shaking at 250 rpm until the optical density at 600 nm (OD_600_) reached 0.7-0.9. Protein expression was then induced with 1 mM isopropyl-β-D-thiogalactopyranoside (IPTG). After 4 h, cells were harvested by centrifugation at 7,000 rpm for 20 min and stored at −80 °C.

### Purification of α-syn Protein

The cell pellet obtained from 1 L of culture was resuspended in 25 mL of lysis buffer (10 mM Tris, pH 8, 1 mM EDTA, 1 mM PMSF, and half of Sigma Fast protease inhibitor cocktail tablet). After transferring the resuspended cells into a centrifuge tube, lysozyme (Final concentration 0.2 mg/ml) was added, and the mixture was incubated on ice for 20-30 min.

Cells were lysed by sonication at pulse rate of 10 s of on-time and 15 s of off-time, with 40% amplitude power for a total on-time of 3.5 min using a Branson Digital Sonifier SFX 250. The lysate incubated on ice for 10-15 min, followed by centrifugation at 16,000 rpm for 1 h at 4 °C.

To maximize purity and yield, an acid precipitation method was adapted^57^. The milky supernatant was transferred to clean 25 mL centrifuge tubes, and the pH was adjusted to 3.5. After stirring at room temperature for 20 min, sample were centrifuged at 16,000 rpm for 30 min at 4 °C, resulting in a clear supernatant. The pH was then adjusted back to 7.5 before an additional centrifugation step at 16,000 rpm for 1 h at 4 °C to remove residual aggregates.

The collected supernatant was filtered through a PVDF syringe filter with 0.45 μm pore size (Avantar, VWR® Syringe Filters) and dialyzed overnight at 4 °C against 1 L of low-salt anionic exchange buffer (10 mM Tris, pH 7.6, 25 mM NaCl, 1 mM EDTA) using a dialysis membrane with a molecular weight cutoff (MWCO) of 3.5 kDa (Spectrum Laboratories, Inc., Spectra/Por®3 Dialysis Membrane).

The following day, the dialyzed α-syn was filtered through a PES syringe filter with 0.22 μm pore size (Avantar, VWR® Syringe Filters) and loaded onto a double-stacked pre-packed HiTrap^TM^ Q Sepharose High Performance anion exchange chromatography column (Cytiva Sweden AB, #17115401) and washed with Buffer A (10 mM Tris, pH 7.6, 25 mM NaCl, 1 mM EDTA) and Buffer B (10 mM Tris, pH 7.6, 1 M NaCl, 1 mM EDTA). α-Syn-containing fractions were pooled at conductivity of 30 mS/cm and concentrated to approximately 13 mg/ml using a 10 kDa Amicon ®Ultra centrifugal filter device (Millipore).

The concentrated protein was loaded onto a pre-packed HiPrep 16/60 Sephacryl S-200 High Resolution preparative size exclusion chromatography column (Cytiva Sweden AB, #17116601). Size exclusion chromatography was performed in a buffer (10 mM Tris, pH 7.6, 100 mM NaCl) with a total volume of 126 ml. α-Syn protein eluted between 40 ml and 60 ml. All column chromatography was performed using an AKTA^TM^ pure (GE Healthcare Bio-Sciences AB, Sweden). Protein concentration was determined using a NanoDrop ND-1000 UV/Vis Spectrophotometer (Thermo Fisher Scientific), and purity was assessed by SDS-PAGE gel electrophoresis. Purified α-syn was dialyzed against 1 L of distilled water overnight at 4 °C and stored at −80 °C.

### Lipid Vesicle Solution

To mimic physiologically relevant normal and aged neuronal membranes, 1-palmitoyl-2-oleoyl-glycero-3-phosphocholine (POPC), 1,2-dioleoyl-sn-glycero-3-phosphoethanolamine (DOPE), 1,2-dipalmitoyl-sn-glycero-3-phosphocholine (DPPC), 1-palmitoyl-2-oleoyl-sn-glycero-3-phosphoethanolamine (POPE), cholesterol, and sphingomyelin (SM) were purchased from Avanti Polar Lipids.

The Neuron membrane model consisted of POPC:DOPE:cholesterol:SM at a molar ratio of 35:20:35:10, while the Aged membrane model consisted of DPPC:POPE:cholesterol:SM at the same molar ratio. These compositions were selected to reflect the lipid abundance of normal neuronal membranes as well as age-associated changes in fatty acid saturation^40,50,51,53^.

To prepare these membranes, phospholipid, cholesterol, and SM were co-dissolved in a chloroform/methanol mixture, and solvents were evaporated under a stream of N_2_ gas. Lipid films were vacuum-desiccated overnight, rehydrated in 12 mM Tris-HCl buffer (pH 7.6) to 12.5 mM total lipid, and briefly sonicated for 10-15 sec. Samples then underwent ten freeze-thaw cycles using liquid nitrogen and a 50 °C water bath to produce homogeneous vesicles. The resulting 12.5 mM vesicle stock was diluted to achieve the desired lipid-to-protein molar (L/P) ratio.

### Fibril and PFFs Formation

Fibrils were generated by incubating 200 μM α-syn in 10 mM Tris-HCl buffer (pH 7.6) for four weeks at 37 °C without agitation, either in the absence or presence of lipid vesicle solutions at L/P ratios of 5, 10, and 50.

Preformed fibrils (PFFs) were generated by sonicating fibrils prepared at L/P = 0 or 10 using a sonifier (Branson Digital Sonifier SFX 250 with Sonifier Sound Enclosure) at a pulse rate of 1s of on-time and 4 s of off-time, 10% amplitude power, and total on-time of 2 min. The average PFF length (∼100 nm) was confirmed by negatively stained transmission electron microscopy (**Fig. S9**). PFFs generated from α-syn fibrils grown in the absence of membranes or in the presence of Neuron or Aged membranes are called PFFs, N-PFFs, Aged-PFFs, respectively. All prepared PFFs were stored at −80 °C and thawed and bath-sonicated for 1 min immediately before use.

To control for lipid-associated contributions in N-PFFs and Aged-PFFs, lipid-free PFFs were mixed with Neuron or Aged membranes at an L/P ratio of 10, generating PFFs + N-lipid and PFFs + Aged-lipid. In parallel, α-syn monomers were combined with Neuron- or Aged membranes (Mono + N-lipid and Mono + Aged-lipid).

### Transmission Electron Microscopy (TEM)

Negatively stained fibrils or PFFs were prepared on lacey carbon-coated copper grids (Electron Microscopy Sciences LC-325-CU-CC, Ted Pella 01824). A 10 μl aliquot was deposited onto the grid and allowed to adsorb for 1-2 min. After absorption, the solution was blotted, washed with 10 μl of water, blotted again, and stained with 10 μl of UranyLess EM stain solution (Electron Microscopy Sciences) for 30 s. TEM images were acquired on a JEOL 2100F Field-emission transmission electron microscope at 120 kV using a Gatan Ultrascan CCD camera and DigitalMicrograph (GMS3) software (Gatan Inc.).

### Thioflavin T (ThT) Fluorescence Monitoring of Fibril Formation Kinetics

ThT (25 μM) was added to a freshly prepared solution of monomeric α-syn (200 μM) in the absence or presence of Neuron or Aged membranes in 10 mM Tris-HCl buffer (pH 7.6). Measurements were performed in Greiner Bio-One 96-well microplates (655900). To prevent sample dehydration during measurements, 100 μl of distilled water was added to the outermost wells.

Fluorescence measurements were recorded on a FLUOstar Omega microplate reader (BMG LABTECH) at 37 °C with excitation at 485 nm and emission at 538 nm. Readings were collected every 30 min or 1 h with a gain setting of 800-1000. Before each measurement, the well plate was orbitally shaken at 700 rpm for 60 s. Each condition was measured in triplicate, and curves were normalized to the maximum fluorescence intensity.

### Solid-State NMR (ssNMR) Sample Preparation

For ssNMR measurements, uniformly ^13^C, ^15^N-labeled α-syn protein was expressed in *E. coli* BL21 (DE3) competent cells. Cells were grown in 2 L of LB media containing ampicillin (100 µg/ml) at 37 °C with shaking at 250 rpm until OD_600_ reached 0.5-0.6. Cells were harvested by centrifugation at 7,000 rpm for 20 min, and the pellet was resuspended in 1 L of M9 medium containing 2 g of ^13^C-labeled glucose, 1 g of ^15^N-labeled ammonium chloride, 2 mM of MgSO_4_, 0.1 mM of CaCl_2_, 100 µg/ml of ampicillin, and 10 mL of Gibco^TM^ 100x MEM vitamin solution (ThermoFisher Scientific). Incubation at 37 °C continued until OD_600_ = 0.8-0.9, at which point expression was induced with 1 mM IPTG. After 4 h, cells were collected by centrifugation at 7,000 rpm for 20 min and stored at −80 °C. Uniformly labeled ^13^C and ^15^N-labeled α-syn was purified using the same procedures as for unlabeled α-syn.

Uniformly ^13^C, ^15^N-labeled lipid-free α-syn fibrils were prepared by seeding 200 µM monomeric ^13^C, ^15^N-labeled α-syn with 5 mol% (monomer equivalent) lipid-free PFFs in 10 mM Tris-HCl buffer (pH 7.6). Uniformly ^13^C, ^15^N-labeled α-syn fibrils grown with Neuron or Aged membranes at an L/P ratio of 10 were prepared using the same procedure as for lipid-free fibrils, except that 5 mol% (monomer equivalent) N-PFFs or Aged-PFFs was added to 200 µM monomeric ^13^C, ^15^N-labeled α-syn in the presence of Neuron or Aged membranes (L/P = 10) in 10 mM Tris-HCl buffer (pH 7.6), respectively. In addition, uniformly ^13^C, ^15^N-labeled α-syn fibrils formed with Aged membranes at an L/P ratio of 50 were prepared by incubating 200 µM monomeric ^13^C, ^15^N-labeled α-syn with Aged membranes (L/P = 50) in 10 mM Tris-HCl buffer (pH 7.6) without seeding. All fibrils used for ssNMR experiments were generated at 37 °C without agitation, and fibril formation was confirmed by negatively stained TEM images.

To evaluate the membrane association of lipid-free α-syn fibrils, uniformly ^13^C, ^15^N-labeled lipid-free α-syn fibrils were mixed with Neuron or Aged membranes at an L/P ratio of 10 and incubated for 24 h at 37 °C without agitation. This enabled direct comparison of membrane association between the two membrane types.

Fibrils were pelleted by centrifugation at 50,000 rpm overnight at 20 °C using a Beckman SW 60Ti rotor. The pellets were slowly dried in a desiccator to achieve 55-65% water by mass. Approximately 25-30 mg of hydrated fibrils were packed into 3.2 mm MAS rotors.

### Solid-State NMR Experiments

All ssNMR experiments were conducted on an 800 MHz (18.8 Tesla) Bruker Advance III-HD NMR spectrometer equipped with a 3.2-mm ^1^H-^13^C-^15^N magic angle spinning (MAS) probe (Black Fox, LLC). ^13^C chemical shifts are reported on the TMS scale using the 38.48 ppm of CH_2_ signal of adamantane as a reference. 1D ^13^C cross-polarization (CP) and 2D ^15^N-^13^C and ^13^C-^13^C correlation spectra were measured under 13.5 kHz MAS at 275-295 K. 2D ^13^C-^13^C DARR spectra were acquired with mixing times of 50 ms and 100 ms^58,59^.

To probe the extent of fibril-membrane association, 2D ^13^C-detected ^1^H-^1^H spin-diffusion experiments were performed on α-syn fibrils grown with Neuron or Aged membranes at 285-295 K under 13.5 kHz MAS^60–62^. A 400 µs ^1^H T_2_ filter was applied to suppress ^1^H magnetization of the rigid fibrils. Following ^1^H chemical shift evolution, ^1^H-^1^H spin diffusion mixing times ranging from 4 ms to 225 ms were used to transfer water and lipid ^1^H magnetization to the protein. The transferred magnetization was detected via ^13^C following ^1^H-^13^C CP.

To ensure that sample integrity was maintained throughout the ssNMR experiments, 1D ^1^H and ^13^C spectra were recorded before and after 2D ssNMR measurements.

### Calcein Release Assays

The ability of α-syn to disrupt the membrane integrity of lipid vesicles was assessed using calcein release assays, following a previously established protocol^63^. To prepare calcein-loaded lipid vesicles, freshly prepared lipid films with the normal neuronal membrane-mimetic composition (POPC:DOPE:cholesterol:SM at a molar ratio of 35:20:35:10) were hydrated with 1 mL of 100 mM calcein solution (pH 7.4) and subjected to 10 freeze-thaw cycles to facilitate calcein encapsulation. The resulting solution was extruded using a mini-extruder (Avanti Polar Lipids) through a 100 nm pore-size polycarbonate membrane (Cytiva) 15 times to generate a homogeneous population of unilamellar liposomes.

Calcein-loaded vesicles were then separated using a gravity-flow chromatography column packed with Sephadex G-50 beads. Eluted fractions containing calcein liposomes, identified by a light orange color, were collected, while darker orange fractions corresponding to free calcein dye were discarded. The purified calcein-containing liposomes were stored at 4 °C until use.

Fluorescence measurements were recorded every 10 min overnight using a FLUOstar Omega microplate reader (BMG LABTECH) with an excitation at 485 nm and an emission at 520 nm. All experiments were conducted at room temperature in Greiner Bio-One 96-well microplates (655900), with a total sample volume of 75 μl per well.

Membrane permeability was assessed by adding 13 μM of α-syn PFFs to purified calcein-containing liposomes. As a positive control for maximum dye release, calcein-containing liposomes were treated with 10% Triton X-100. Membrane permeability was calculated using the following equation:

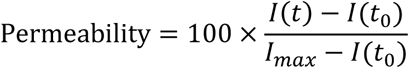

where 𝐼(𝑡) represents the fluorescence intensity at time t, 𝐼(𝑡_0_) is the initial fluorescence intensity at t = 0, and 𝐼_𝑚𝑎𝑥_ is the maximum fluorescence intensity measured after treatment with 10% Triton X-100.

### Bicinchoninic Acid (BCA) Assay for Monomer-Membrane Binding Analysis

The concentration of membrane-bound α-syn monomers was quantified using a bicinchoninic acid (BCA) assay. Freshly prepared monomeric α-syn (200 µM), with or without Neuron or Aged membranes (L/P = 10) in 10 mM Tris-HCl buffer (pH 7.6), was vortexed 5 s and incubated quiescently at 37 °C for 4 h to allow α-syn-membrane binding. Following incubation, membrane-bound α-syn was pelleted by centrifugation at 50,000 rpm for 1 h at 4 °C.

A 20 μL of the supernatant containing unbound α-syn monomer was carefully collected for denaturation and BCA analysis. Control samples consisting of 10 mM Tris-HCl buffer, 2 mM Neuron membranes, or 2 mM Aged membranes without α-syn monomers were processed in parallel.

A six-point protein standard curve (0-200 µM α-syn monomer) was prepared in 10 mM Tris-HCl buffer (pH 7.6). A denaturing solution (0.2 M NaOH + 2% SDS) was prepared in deionized water. For each standard or sample, 20 μL of solution was mixed with 20 μL denaturing solution, vortexed briefly, and heated at 95 °C for 5 min to fully denature proteins and then cooled to room temperature.

BCA working reagent (WR) was freshly prepared by mixing BCA Reagent A and Reagent B (ThermoFisher Scientific) at a 50:1 (A:B) ratio. For each reaction, 25 μL of the denatured sample or standard was added to 200 μL of BCA WR, vortexed for 5 s, and incubated at 37 °C for 30 min. After incubation, 50 μL aliquots of each reaction were dispensed into a Greiner Bio-One 96-well microplate (655900) in quadruplicate. Absorbance at 562 nm was measured using a FLUOstar Omega microplate reader (BMG LABTECH). Readings were collected every 10 min for 40 min, with orbital shaking at 700 rpm for 60 s before each measurement. The average absorbance from four measurements was reported.

### Dynamic Light Scattering (DLS) Measurements

DLS measurements were performed using a Zetasizer Nano Series Instrument (Malvern Instruments) equipped with a He-Ne laser (λ = 633 nm) and operated at a backscattering detection angle of 173°. Neuron or Aged membrane samples were prepared as described above using 10 freeze-thaw cycles, then diluted to 2 mM lipid concentration either in the absence or presence of 200 µM α-syn monomers in 10 mM Tris-HCl buffer (pH 7.6). Samples were incubated at 37 °C for 0, 2, 7, and 14 days prior to measurements.

Before measurement, samples were diluted 1:1 (v/v) with 10 mM Tris-HCl buffer (pH 7.6), then resuspended and mixed thoroughly before transfer into a disposable quartz cuvette (ZEN0040, Malvern Panalytical). All measurements were conducted at 25 °C using water as the dispersant (refractive index = 1.330; viscosity = 0.8872 cP at 25 °C). The sample viscosity was set equal to that of the dispersant. Phospholipid optical parameters were set to a refractive index of 1.450 and absorption coefficient of 0.100. For each condition, 15 runs of 10 s were recorded following a 2 min equilibrium period. Data were analyzed using Zetasizer software (Malvern Instruments) with the general-purpose analysis model.

### Proteinase K Digestion

Proteinase K (PK) digestion was performed on three types of 5 mol% seeded α-syn fibril prepared in 10 mM Tris-HCl buffer (pH 7.6): lipid-free α-syn fibrils, α-syn + Neuron fibrils, and α-syn + Aged fibrils. Mature fibrils were mixed with PK to a final PK concentration of 0.05 mg/mL. After a brief 5-10 s vortex, samples were incubated at 37 °C for 2-10 min. Reactions were terminated by heating samples at 90 °C for 15 min. Samples were then cooled to room temperature and briefly centrifuged. PK-inactivated samples were snap-frozen in liquid nitrogen, lyophilized, and resuspended in 200 μL of hexafluoroisopropanol (HFIP). Samples were sonicated in a water bath for 5 min, vortexed for 30 s, and left capped in a fume hood overnight to allow complete HFIP solubilization. The following day, HIFP was evaporated under a gentle stream of N_2_ gas.

To remove lipid components, an acetone precipitation step was performed. Pre-cooled acetone (−20 °C) was added at four times the sample volume, followed by vortexing for 10 s and incubated at −20 °C for 1 h. Samples were centrifuged at 13,200 rpm for 10 min, the supernatant was discarded, and residual acetone was evaporated under N_2_. Dried pellets were resuspended in 50 μL of denaturing buffer (8 M urea and 4% SDS), vortexed vigorously, sonicated for 1 min, and incubated at room temperature for 15 min with intermittent vortexing to ensure complete solubilization.

Protein digestion profiles were analyzed by SDS-PAGE followed by silver staining. Gels were fixed twice for 15 min in 30% ethanol:10% acetic acid, washed sequentially with 10% ethanol and distilled water, sensitized for 1 min, stained for 30 min, and developed for 30 s to 2 min until optimal band intensity was achieved. Development was stopped with 5% acetic acid for 10 min.

### Cell Culture

To establish a dopaminergic neuron model, the human neuroblastoma cell line (SH-SY5Y; ATCC, CRL-2266) was selected and expanded in 25 cm^2^ flasks following the manufacturer’s protocols. To culture SH-SY5Y cells, Dulbecco’s Modified Eagle Medium (DMEM; 10-017-CV, Corning) was used as the basal growth medium, supplemented with 10% (v/v) heat-inactivated fetal bovine serum (hiFBS; SH30396.03, Cytiva), 1% (v/v) GlutaMAX-I (35050-06, ThermoFisher) and 1% (v/v) penicillin-streptomycin (SV30010, Cytiva). SH-SY5Y cells at passages 3 to 10 were used as recommended by the cell supplier.

The process of dopaminergic neuronal differentiation started with trypsinization and resuspension of SH-SY5Y cells in growth medium. The harvested cells were then introduced into 48-well plates at a density of 5.0 × 10^5^ cells/mL. The initial culture was maintained in their growth medium to promote cell adherence and stabilization. Two days after seeding, the SH-SY5Y cell growth medium was replaced with the first differentiation medium consisting of DMEM supplemented with 2.5% (v/v) hiFBS, 1% (v/v) GlutaMAX-I, 1% (v/v) penicillin-streptomycin, and 10 μM retinoic acid (50-165-6969, Fisher Scientific) to induce differentiation into dopaminergic neuronal cells. After 5 days, the first differentiation medium was switched to a second differentiation medium based on Neurobasal-A (10888022, ThermoFisher) mixed with 50 ng/mL brain-derived neurotrophic factor (BDNF; P23560, FUJIFILM Irvine Scientific), 20mM potassium chloride (KCl; 7447-40-7, Avantor Science), 1% (v/v) B27 (17-504-044, Fisher Scientific), 1% (v/v) GlutaMAX-I, and 1% (v/v) penicillin-streptomycin. The second differentiation medium was also maintained for an additional 5 days. Both the first and second differentiation media were refreshed every other day to support and sustain the dopaminergic differentiation of SH-SY5Y cells.

### Treatment of Lipid-free or Membrane-associated α-syn PFFs

To investigate the pathological impacts of polymorphic α-syn fibrils, differentiated dopaminergic neuronal cells were exposed to different α-syn PFFs. A 200 µM stock solution of each α-syn PFFs conformer was sonicated for 15 s to ensure uniform dispersion and achieve an average fibril length of approximately ∼100 nm. For cell treatment, the sonicated PFF stock solution was diluted to a final concentration of 1 μg/mL in the second differentiation medium.

On Day 12 of culture, α-syn PFFs-containing medium was added to each well containing dopaminergic neuronal cells. The cultures were then incubated with the fibrils for 72 h. After incubation, the medium was replaced with fresh differentiation medium. Treated cells were either analyzed immediately or maintained for an additional 3 days prior to analysis.

### Immunostaining and Quantification

For immunostaining, dopaminergic neuronal cells were fixed with 4% paraformaldehyde (AR1068, Boster) for 30 min at RT and washed twice using DPBS. After fixation, cells were permeabilized with 0.1% Triton X-100 (9002-93-1, VWR) for 15 min. To prevent non-specific binding, the cells were blocked with 1.5% bovine serum albumin (BSA; 22013, Biotium) for 30 min at RT. Subsequently, the cells were incubated overnight at 4°C with primary antibodies that selectively bind to the proteins of interest.

To visualize the actin cytoskeleton, Phalloidin-CF 430 (Biotium) was applied at 1:100 dilution. Neuronal differentiation was assessed using primary antibodies: rabbit polyclonal anti-MAP2 (A16829, 1:200, Antibodies.com) for dendritic extensions, mouse monoclonal anti-GAP43 (A85392, 1:1000, Antibodies.com) for axonal projections, and mouse monoclonal anti-β III Tubulin (A86691, 1:500, Antibodies.com) for the neuronal cytoskeleton. To confirm PD-specific misfolding of endogenous α-syn and α-syn aggregation, we used rabbit monoclonal antibodies against α-syn phosphorylated at Ser129 (A304933, 1:500, Antibodies.com) and α-syn aggregates (A209538, 1:5000, Antibodies.com), respectively. To assess the dopaminergic characteristics of neuronal cells, Tyrosine Hydroxylase (TH) expression was examined using a mouse monoclonal anti-TH antibody (A104316, 1:5000, Antibodies.com). For investigating inflammatory responses, a rabbit polyclonal anti-NF-kB antibody (51-0500, 1:500, ThermoFisher) was used.

After incubation with primary antibodies, cells were washed twice with DPBS and incubated for 1-2 hours at RT with fluorescently labeled secondary antibodies (A32732, ThermoFisher; A32723, ThermoFisher; A32731, ThermoFisher; A11032, ThermoFisher). Hoechst (33342, ThermoFisher) was used for nuclear staining.

Fluorescence images of the cells were captured using a Nikon Ti2-Eclipse microscope. NF-κB and pSer129 imaging was performed with excitation and emission wavelengths of 500 nm and 535 nm, respectively, and α-syn fibrils labeled with MJFR-14-6-4-2 were imaged with excitation at 620 nm and emission at 700 nm. Exposure times were 900 ms for NF-κB, pSer129, and TH, and 300 ms for MJFR-14-6-4-2. Image processing was performed using NIS Elements software.

To quantitively assess the data, fluorescence intensity was averaged from three samples for each experimental condition. To account for background fluorescence due to light diffusion, background signal was measured in acellular regions and subtracted from cellular measurements, thereby correcting for diffuse light that uniformly elevates fluorescence intensity.

Statistical value of the obtained data was assessed by a two-tailed t-test. Results from three independent experiments are presented as mean ± SD.

## RESULTS

### α-syn Forms Fibrils in the Presence of Normal and Aged Neuronal Membranes

To model physiological conditions of neuronal membranes and their age-related alterations, we prepared two types of vesicles: Neuron membranes, representing normal neuronal plasma membranes, and Aged neuronal membranes. Neuron membranes were composed of 1-palmitoyl-2-oleoyl-glycero-3-phosphocholine (POPC), 1,2-dioleoyl-sn-glycero-3-phosphoethanolamine (DOPE), cholesterol, and sphingomyelin (SM) at a 35:20:35:10 molar ratio. To reflect the reduction in unsaturated fatty acid content and acyl chain length observed in brains with aging, Aged membranes were prepared by substituting POPC and DOPE with 1,2-dipalmitoyl-sn-glycero-3-phosphocholine (DPPC) and 1-palmitoyl-2-oleoyl-sn-glycero-3-phosphoethanolamine (POPE), while maintaining the same molar ratios (**Fig. 1**). α-syn fibril formation was initiated by incubating 200 μM of α-syn monomers at 37 °C under quiescent conditions either in the absence or presence of Neuron or Aged membranes at a lipid-to-protein molar (L/P) ratio of 10. Structural changes in α-syn monomers during incubation were monitored by circular dichroism (CD) spectroscopy (**Fig. 2D**). Initially, α-syn monomers incubated with Neuron (green) or Aged (red) membranes exhibited α-helical-like spectra, characterized by two minima near 208 and 222 nm, consistent with previous reports showing that membrane-bound α-syn adopts a helical structure^64–66^. After 400 h of incubation without agitation, a pronounced negative peak appeared near 218 nm, indicating a transition to a β-sheet-rich conformation, a hallmark of fibril formation. These results confirm that α-syn monomers form fibrils over extended incubation periods. Fibril formation, in both the absence and presence of membranes, was also confirmed by negatively stained transmission electron microscopy (TEM) (**Fig. 2A-C**).

**Figure 1.**
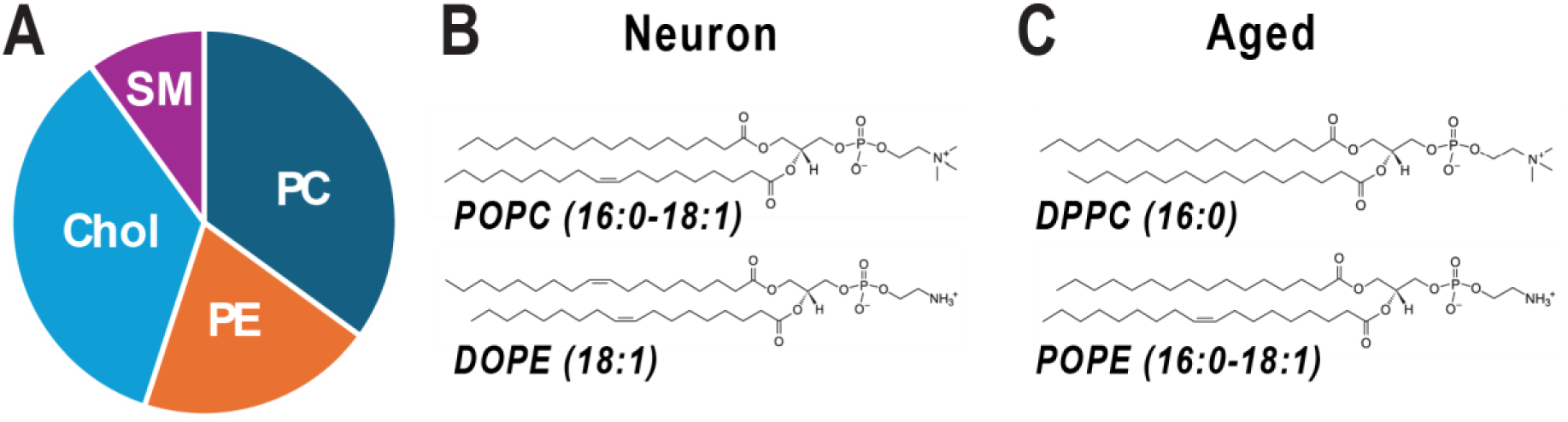
Lipid compositions of model membranes mimicking plasma membranes of normal and aged neurons, referred to as Neuron and Aged membranes in this study. (A) Lipid compositions of Neuron and Aged membranes, consisting of PC, PE, Cholesterol, and SM in a 35:20:35:10 molar ratio. (B) Chemical structures of POPC and DOPE, the primarily lipid components of the Neuron membrane. (C) Chemical structures of DPPC and POPE, which characterize the Aged membrane with altered saturation levels. Compared to POPC and DOPE in Neuron membranes, DPPC and POPE have shorter and more saturated acyl chains.

**Figure 2.**
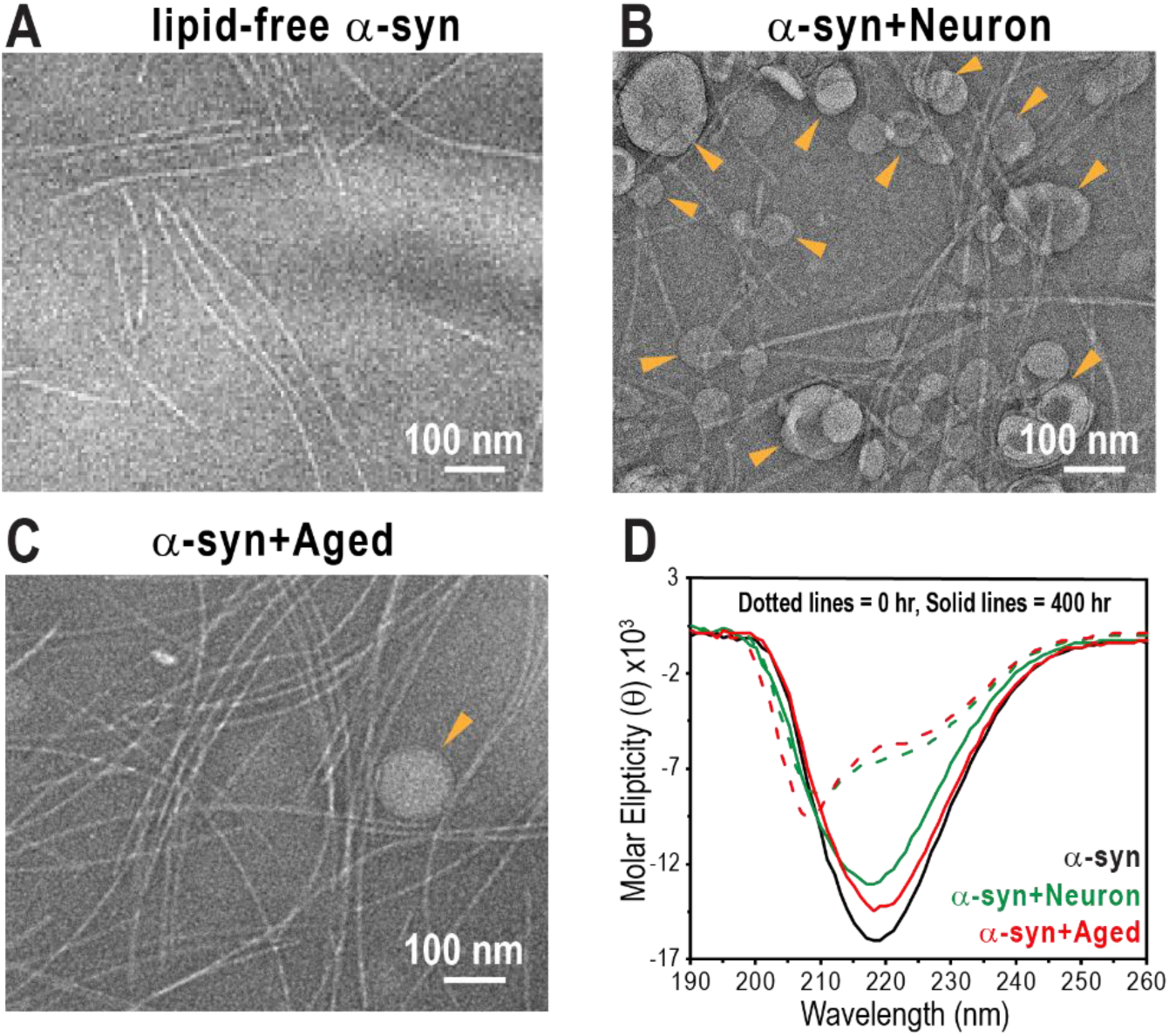
Characterization of α-syn fibril formation in the absence and presence of lipid membranes. (A-C) Negatively stained TEM images confirming fibril formation after incubation: (A) α-syn fibrils formed without membranes, (B) α-syn fibrils formed in the presence of Neuron membranes, and (C) α-syn fibrils formed in the presence of Aged membranes. Scale bars: 100 nm. Orange arrows indicate lipid membranes. (D) CD spectra of α-syn in the absence (black) and presence of Neuron (green) or Aged membrane (red) over 400 hours of incubation without agitation. Dotted lines represent spectra at 0 hours, and solid lines indicate spectra at 400 hours, revealing structural transitions from an α-helical conformation to a β-sheet-rich fibril structure.

### Membranes Alters a-syn Aggregation Kinetics

Thioflavin T (ThT) fluorescence assays were used to monitor aggregation kinetics under varying protein concentrations and L/P ratios, with three replicates measured for each condition. Each fluorescence curve was normalized to its maximum intensity (**Fig. 3A-C**). The t_0.1_ and t_0.5_ values—corresponding to the times required to reach 10% and 50% of maximal ThT fluorescence intensity, respectively—were used to quantitatively compare aggregation properties under different conditions (**Fig. 3D**). These parameters are commonly referred to as the lag phase and elongation phase, reflecting monomer nucleation and subsequent conversion into amyloid forms, respectively^20^. In the absence of lipids, accelerated fibril formation was observed at higher α-syn monomer concentrations, as evidenced by decreased t_0.1_ and t_0.5_ values (**Fig. 3D**). To assess the effects of Neuron and Aged membranes on fibril formation, all subsequent experiments were conducted at a monomer concentration of 200 μM with L/P ratios of 5, 10, and 50 (**Fig. 3B, C**).

**Figure 3.**
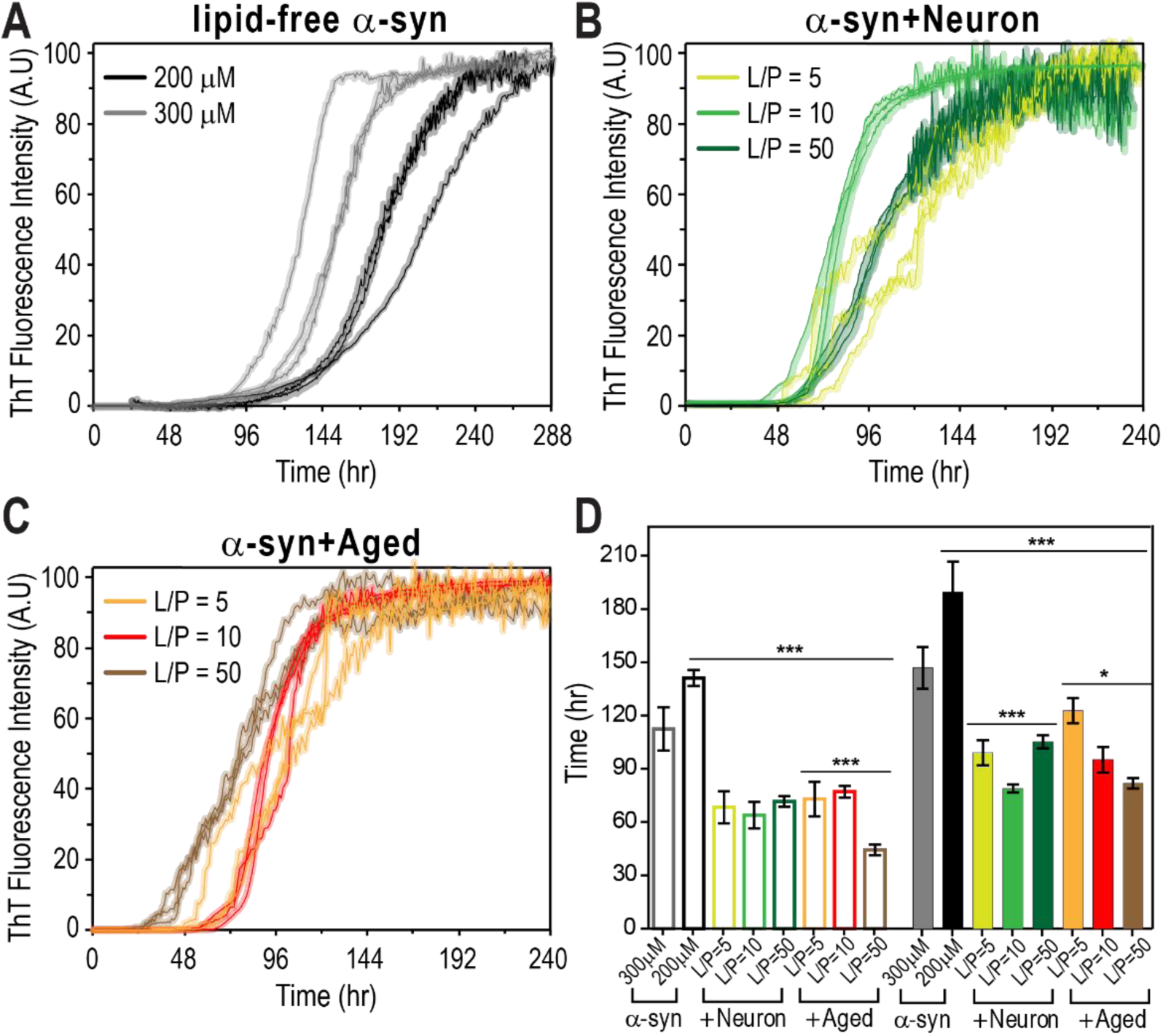
Effects of protein concentration and L/P ratio on α-syn aggregation kinetics. (A) ThT fluorescence curves of α-syn aggregation kinetics at monomer concentration of 200 μM (black) and 300 μM (gray) in the absence of lipid over 288 hours. Higher α-syn concentrations result in shorter lag times and steeper sigmoidal curves. (B) ThT fluorescence curves of α-syn aggregation in the presence of Neuron membranes at L/P ratios of 5 (lime green), 10 (green), and 50 (dark green) over 240 hours. (C) ThT fluorescence curves of α-syn aggregation in the presence of Aged membranes at L/P ratios of 5 (orange), 10 (red), and 50 (brown) over 240 hours. (D) Quantitative comparison of aggregation kinetics using t_0.1_ (open bars) and t_0.5_ (solid bars), representing the times required to reach 10% and 50% of maximal ThT fluorescence intensity, respectively. The same color scheme from (A-C) is applied. Data show mean ± SD with n = 3. ***P < 0.001, **P < 0.01 *P < 0.05.

The presence of both Neuron and Aged membranes enhanced α-syn fibril formation at 200 µM across all L/P ratios, as indicated by significantly reduced t_0.1_ and t_0.5_ values. For Neuron membranes (**Fig. 3B**), the data suggest that an L/P ratio of 10 yields a shorter elongation phase compared to L/P ratios of 5 and 50, accompanied by steeper sigmoidal ThT curves. We attribute this effect to membrane surface-bound α-syn, which acts as a nucleation site, promoting aggregation at relatively low L/P ratios. However, at an L/P ratio of 50, increased α-syn binding to lipid vesicles reduces the concentration of free α-syn monomers in solution, thereby slowing fibril formation. Consistently, α-syn monomer binding assays showed higher levels of membrane-bound α-syn monomers with Neuron membranes at an L/P ratio of 50 compared to L/P ratios of 5 and 10 (**Fig. S6A**).

In contrast, in the presence of Aged membranes (**Fig. 3C**), an overall trend toward faster fibril formation was observed with higher L/P ratios. Notably, at an L/P ratio of 50, both the lag phase a^67^nd the elongation phase were markedly shortened (**Fig. 3D**). This observation suggests that α-syn has a lower binding affinity for Aged membranes than for Neuron membranes, allowing a higher concentration of free α-syn to remain in solution and thereby facilitating fibril formation at high L/P ratios. Consistent with this interpretation, the lower levels of membrane-bound α-syn monomers observed with Aged membranes at L/P ratios of 5 and 50, further support this conclusion (**Fig. S6A**).

### Membrane-Induced Conformational Variations in α-syn Fibrils Revealed by Solid-State NMR

While the TEM images of the three α-syn fibril samples primarily show untwisted fibrils (**Fig. 2A-C**), minor populations of twisted fibrils with distinct crossover patterns were also observed (**Fig. S1)**. These observations indicate conformational polymorphism both within and between samples, likely arising from the complex mechanisms underlying membrane-associated protein aggregation.

Protein aggregates typically consist of a protease-resistant fibrillar core surrounded by a protease-susceptible disordered region. Accordingly, proteinase K (PK) digestion is commonly used to compare structural differences among fibril strains^67–70^. To assess conformational variations among three α-syn fibrils grown either in the absence or in the presence of Neuron or Aged membranes, the fibrils were treated with PK, and the resulting digestion patterns were analyzed by SDS-PAGE (**Fig. S2**). α-Syn fibrils formed in the presence of Neuron membranes exhibit a protease-resistant core after 2 min of proteinase K (PK) digestion but are largely degraded after 10 min, as indicated by the appearance of fragments smaller than ∼11 kDa. In contrast, lipid-free fibrils and fibrils formed with Aged membranes retain PK-resistant cores even after 10 min of digestion. Notably, these two samples display distinct digestion patterns, differing in both the sizes and relative intensities of the resulting peptide fragments. Together, these results demonstrate that all three fibril preparations possess structurally distinct cores.

We next measured 2D ssNMR spectra for uniformly ^13^C, ^15^N-labeled α-syn fibrils formed either without lipids or in the presence of Neuron or Aged membranes at an L/P ratio of 10 for detailed structural comparison (**Fig. 4A-C**). Fibril growth for lipid-free α-syn, α-syn + Neuron, and α-syn + Aged membrane samples was initiated by adding 5 mol% (monomer equivalent) of the corresponding short, preformed fibrils (PFFs). Specifically, three types of PFFs were generated by sonicating fibrils grown without lipids, with Neuron, or with Aged membranes, referred to as PFFs, N-PFFs, Aged-PFFs, respectively. The resulting PFFs were approximately 100 nm in length, as confirmed by negatively stained TEM (**Fig. S9**). Each PFF type was then added to 200 µM uniformly ^13^C, ^15^N-labeled α-syn monomers in the absence or presence of the corresponding membranes (L/P = 10).

**Figure 4.**
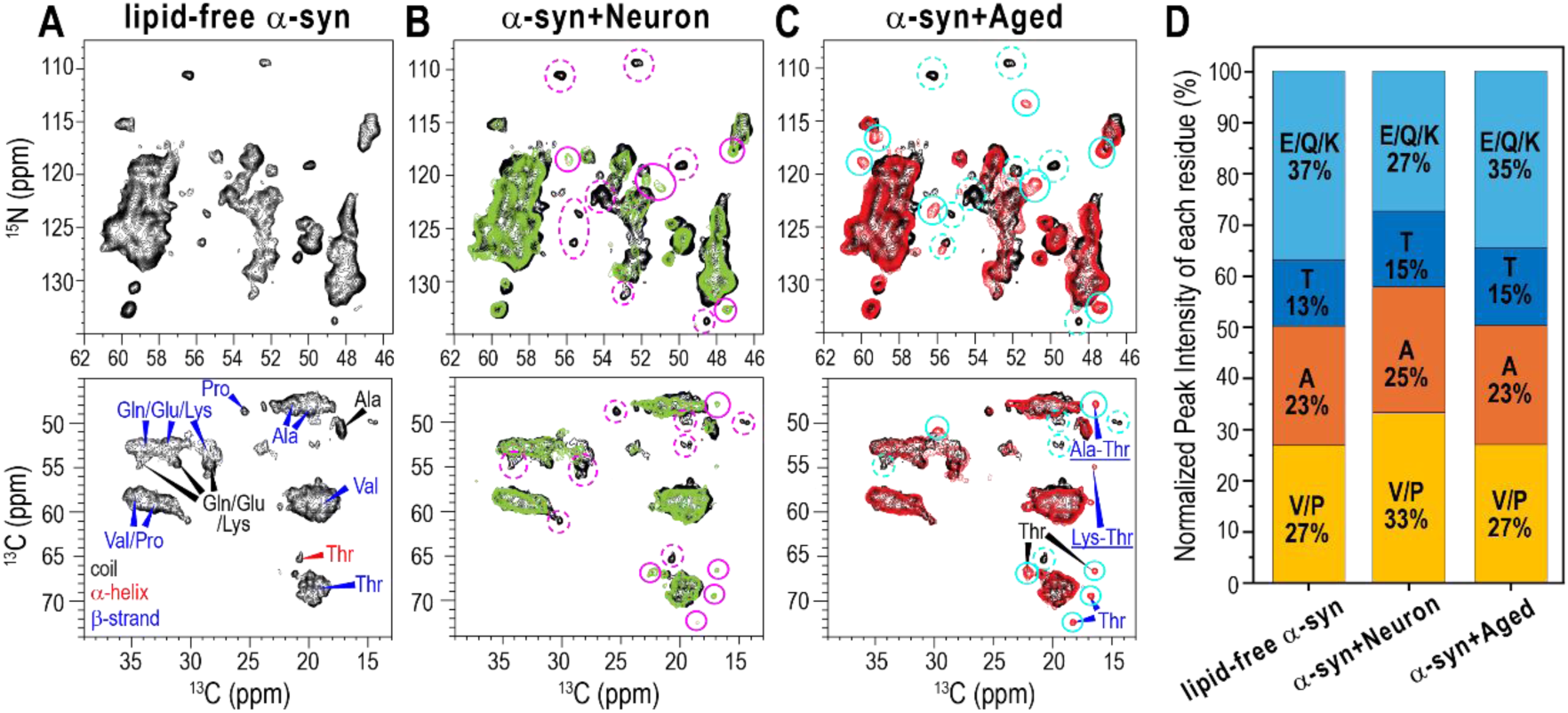
Comparison of 2D NMR spectra of α-syn fibrils grown under different conditions. (A) 2D ^15^N-^13^C (top) and ^13^C-^13^C (bottom) correlation spectra of lipid-free α-syn fibrils (black). (B) 2D ^15^N-^13^C (top) and ^13^C-^13^C correlation spectra of α-syn fibrils grown with Neuron membranes (green), overlaid with lipid-free α-syn fibrils (black). Dotted pink circles indicate peaks present only in lipid-free α-syn fibrils, while solid pink circles highlight peaks unique to α-syn fibrils grown with Neuron membranes. (C) 2D ^15^N-^13^C (top) and ^13^C-^13^C (bottom) correlation spectra of α-syn fibrils formed with Aged membranes (red), overlaid with the lipid-free α-syn fibrils (black). Dotted cyan circles indicate peaks present only in lipid-free α-syn fibrils, while solid cyan circles mark peaks exclusive to α-syn fibrils formed with Aged membranes. (D) Quantification of residue-specific Cα-Cβ cross peak intensities from the 2D ^13^C-^13^C spectra, normalized within each dataset, comparing the relative amino acid composition of fibril core among lipid-free α-syn fibrils, α-syn fibrils grown with Neuron, and those grown with Aged membranes.

All ssNMR spectra were acquired under conditions optimized for ssNMR, using cross-polarization (CP) which transfers nuclear spin polarization through magnetic dipole-dipole couplings^71^, along with high-power ^1^H decoupling^72^. Under these conditions, ssNMR detects signals primarily from rigid, immobilized residues within the fibril core^73–76^.

In the 2D ^15^N-^13^C spectra, all three fibril samples exhibited most peaks with ^15^N chemical shifts exceeding 120 ppm, indicating β-sheet conformations. However, several distinct peaks, highlighted by pink and cyan circles, confirm membrane-induced conformational variations among the three fibrils (**Fig. 4B, C**). To obtain amino acid-specific structural information of the fibril core, we measured 2D ^13^C-^13^C correlation spectra using a dipolar coupling-based polarization transfer sequence. The 2D ^13^C-^13^C correlation spectrum of lipid-free α-syn fibrils, acquired with a 100 ms DARR mixing time, shows β-sheet chemical shifts for Thr, Val/Pro, Gln/Glu/Lys, and Ala residues, along with a small helical Thr peak and random coil peaks for Ala and Gln/Glu/Lys residues. This confirms that lipid-free α-syn fibrils adopt a β-sheet rich amyloid structure with a small, disordered region (**Fig. 4A)**. The 2D ^13^C-^13^C correlation spectra of α-syn fibrils grown with Neuron or Aged membranes show predominant peaks that align closely with those observed in lipid-free fibrils (**Fig. 4B, C**). However, these membrane-associated fibrils lack the α-helical Thr peak observed in lipid-free fibrils and instead exhibit additional β-sheet Thr peaks. Fibrils grown with Neuron membranes show weaker coiled Gln/Glu/Lys peaks compared to the other two fibrils. These observations indicate conformational differences in the fibril core across the three fibril samples. Notably, fibrils grown with Neuron or Aged membranes display inter-residue cross peaks between Thr-Ala or Thr-Lys, despite being measured with a shorter 50 ms DARR mixing time compared to lipid-free fibrils. This observation suggests that membrane-associated fibrils adopt a more compact and rigid core structure. Considering the multiple Thr-Lys and Thr-Ala pairs in the α-syn sequence, these inter-residue correlations likely result from intra-molecular interactions.

Because 2D ^13^C-^13^C correlation spectra capture signals from the rigid fibril core, the measured peak intensities reflect the relative abundance of residues present in the fibril core. All 2D ^13^C-^13^C spectra of α-syn fibrils exhibited well-resolved Cα-Cβ cross peaks for Val/Pro, Ala, Thr, and Gln/Glu/Lys residues, enabling quantitative comparison of their relative abundances. Peak intensities were obtained by integrating each corresponding cross peak and were normalized to the sum of the four peaks to allow comparison of relative contributions within each 2D ^13^C-^13^C correlation spectrum (**Fig. 4D**). For lipid-free α-syn fibrils (**Fig. 4A**), the normalized intensities for Val/Pro, Ala, Thr, Gln/Glu/Lys were 27%:23%:13%:37%. In contrast, α-syn fibrils grown with the Neuron membranes displayed 33%:25%:15%:27%, showing an increased Val/Pro contribution and reduced Gln/Glu/Lys contribution. Fibrils grown with Aged membranes showed ratios of 27%:23%:15%:35%, closely resembling lipid-free α-syn fibrils but with slightly higher Thr and reduced Gln/Glu/Lys content. Consistent variations were also observed in the 1D ^13^C CP spectra of the three fibrils (**Fig. S5I**). In lipid-free α-syn fibrils, the spectrum shows stronger signals for Gln/Glu/Lys residues in the 52-55 ppm range and weaker Thr peaks in the 66-69 ppm range compared to the two membrane-associated fibrils. Together, these results demonstrate distinct amino-acid intensity profiles across the three α-syn fibrils, indicating that α-syn fibrils adopt structurally different fibril core domains under varying membrane conditions.

We next used the normalized peak intensities to estimate the fibril core regions by comparing experimental peak ratios with the amino acid composition of different segments of the α-syn sequence. For lipid-free fibrils, the normalized peak intensities closely matched the amino acid distribution of residues M1-L100, suggesting that the fibril core primarily consists of the N-terminal ∼100 residues of α-syn, while G101-A140 remain flexible and do not contribute to the rigid core (**Fig. S5B, C**). In contrast, α-syn fibrils grown with Neuron membranes closely matched the composition of residues G36-A91, whereas fibrils formed with Aged membranes aligned with residues G14-L100 (**Fig. S5D-G**). While these findings support membrane-dependent variations in the fibril core, ssNMR spectra represent averaged signals from all conformations present in a heterogenous sample. Therefore, additional structural analyses will be necessary to precisely define the fibril core regions, quantitatively assess conformational heterogeneity, and further characterize membrane-induced structural changes.

### Distinct Membrane Contacts of α-syn Fibrils with Neuron and Aged Membranes

To investigate the molecular contacts between α-syn fibrils and membranes, we performed 2D ^13^C-detected ^1^H-^1^H spin diffusion NMR experiments on α-syn fibrils grown with Neuron or Aged membranes at an L/P ratio of 10. This experiment correlates the ^1^H signals from water or lipid acyl chains with ^13^C signals from the protein by selectively transferring water and lipid ^1^H signals to the ^13^C-labeled protein using a 400 μs T_2_ filter, followed by varying ^1^H-^1^H spin diffusion mixing times ranging from 4 to 225 ms. The extent of fibril-membrane and fibril-water interactions can be assessed by comparing the intensities of protein signals transferred from lipids and water. **Fig. 5A** and **5C** show representative 2D ^1^H-^13^C spectra of α-syn fibrils grown with Neuron and Aged membranes, respectively, measured with a ^1^H-^1^H spin diffusion mixing time of 100 ms. In both samples, ^13^C-^1^H cross peaks between α-syn and lipids, arising from magnetization transfer from lipids to protein, are observed, indicating that both fibrils contact lipid membranes. Notably, α-syn fibrils grown with Neuron membranes exhibited stronger lipid-to-protein signal transfer than those grown with Aged membranes, implying stronger membrane interactions in the former.

**Figure 5.**
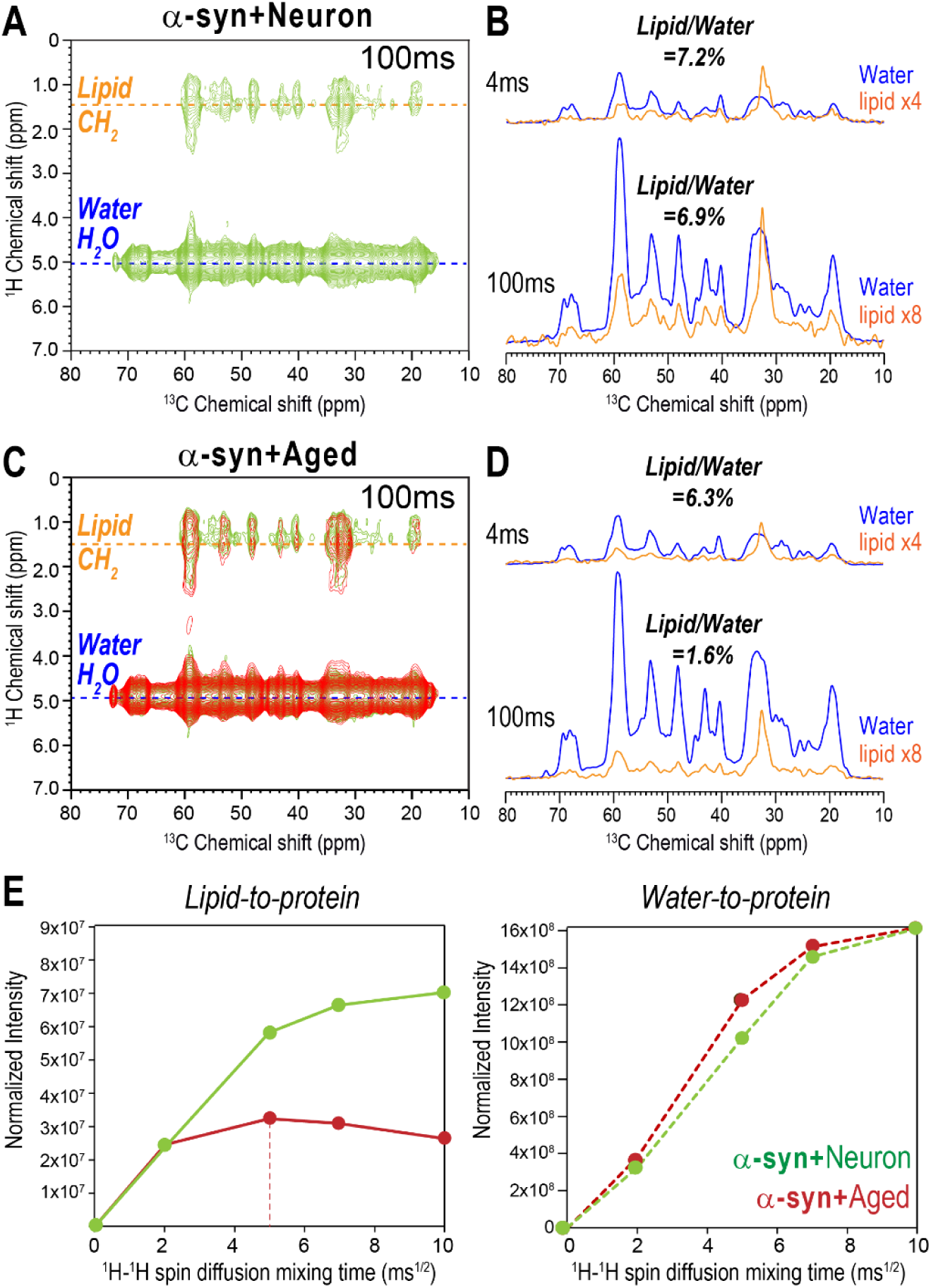
Membrane association of α-syn fibrils with Neuron and Aged membranes. (A, C) Representative 2D ^13^C-detected ^1^H-^1^H spin diffusion NMR spectra of α-syn fibrils grown with Neuron membranes (A, green) and with Aged membranes (C, red), overlaid with spectra of fibrils formed with Neuron membranes (green) at an L/P ratio of 10. All spectra were acquired with a ^1^H-^1^H spin diffusion mixing time of 100 ms. (B, D) ^13^C cross-sections at ^1^H chemical shifts of lipid acyl chain (1.3 ppm, orange) and water (5.0 ppm, blue), extracted from 2D ^1^H-^13^C spectra of α-syn fibrils grown with Neuron (B) and Aged (D) membranes measured at ^1^H-^1^H spin diffusion mixing times of 4 ms and 100 ms. Lipid cross-sections are scaled by a factor of 4 or 8 relative to the water cross-sections for clarity. At 100 ms, α-syn fibrils grown with Neuron membranes exhibit higher lipid-to-protein cross peak intensity (7.2%) than those grown with Aged membranes (6.3%), suggesting stronger membrane interactions with Neuron membranes. (E) Lipid-to-protein (solid lines, left) and water-to-protein (dotted lines, right) ^1^H polarization transfer buildup curves as a function of ^1^H-^1^H spin diffusion mixing time. Protein signals transferred from Aged membranes reach saturation after 25 ms of ^1^H-^1^H spin diffusion mixing time, whereas signals from Neuron membranes (green) continue to increase and remain stronger throughout the mixing-time range.

To quantitatively compare the membrane association of the two fibrils, we extracted 1D cross-sections of ^13^C protein signals transferred from lipid CH_2_ (1.3 ppm) and water (5.0 ppm) in the 2D spectra acquired with ^1^H-^1^H spin diffusion mixing times of 4 ms and 100 ms and compared their intensities. At a 4 ms mixing time, both fibrils show a lipid-to-protein cross peak intensity of approximately 6-7% relative to the water-to-protein cross peak intensities (**Fig. 5B** and **5D**). This ratio is below the typical range observed for membrane-spanning proteins (10-25%)^77–79^, suggesting that neither fibril type is deeply embedded into the membrane bilayer. At a 100 ms of ^1^H-^1^H spin diffusion mixing time, α-syn fibrils grown with Neuron membranes maintain a similar ∼7% lipid-to-protein cross peak intensity relative to water-to-protein cross peaks, indicating that both lipid and water signals increase proportionally with longer ^1^H-^1^H spin diffusion mixing times. In contrast, α-syn fibrils grown with Aged membranes show a significant increase in water-to-protein cross peak intensity but not in lipid-to-protein intensity at a 100 ms ^1^H-^1^H spin diffusion mixing time, reducing the lipid-to-water cross peak intensity ratio to 1.6%. This suggests that water signal transfer dominates over lipid signal transfer at longer mixing times, implying that α-syn fibrils grown with Aged membranes engage in weaker membrane interactions than those grown with Neuron membranes.

**Fig. 5E** presents lipid-to-protein and water-to-protein ^1^H polarization transfer curves for both fibril types as a function of ^1^H-^1^H spin diffusion mixing time. The intensities at each ^1^H-^1^H spin diffusion mixing time were normalized to the maximum water-to-protein intensity at 100 ms mixing time. The resulting plots reveal that protein signals transferred from Aged membranes plateaued after 25 ms of ^1^H-^1^H spin diffusion mixing time, whereas those from Neuron membranes continued to increase. Additionally, the lipid-to-protein signal intensity for α-syn fibrils associated with Aged membrane is significantly lower, approximately three times less, than for fibrils with Neuron membranes. The significantly weak and rapidly saturated lipid-to-protein signals for α-syn fibrils grown with Aged membranes implies that α-syn fibril insertion into the Aged membrane bilayer is limited to a subset of fibrils, or that some broken membrane fragments are incorporated into the fibrils. By contrast, α-syn fibrils grown with Neuron membranes display a steady increase in lipid-to-protein signal intensity up to 225 ms of ^1^H-^1^H spin diffusion mixing time (data not shown), indicating that these fibrils not only insert into the lipid bilayer but also remain associated with surrounding Neuron vesicles, as observed in the TEM image (**Fig. 2B**).

The observed differences in membrane association may stem from stronger binding affinities of Neuron membranes to α-syn monomers or fibrils compared with Aged membranes. Therefore, we measured the binding affinities of α-syn monomers and fibrils to Neuron and Aged membranes. α-syn monomers were incubated with Neuron or Aged membranes for 4 hours at 37 °C, after which membrane-bound monomers were separated by ultracentrifugation, and the concentration of free α-syn monomers remaining in the supernatant was quantified^80,81^. Although α-syn monomers exhibited comparable binding affinities to Neuron and Aged membranes at an L/P ratio of 10, higher monomer binding affinities were observed for Neuron membranes than Aged membranes at L/P ratios of 5 and 50. (**Fig. S6A**). Next, we examined whether Neuron membranes exhibit stronger binding to lipid-free α-syn fibrils than Aged membranes. Lipid-free α-syn fibrils were incubated with Neuron or Aged membranes at an L/P ratio of 10, and membrane-fibril interactions were assessed by conducting ^13^C-detected ^1^H-^1^H spin diffusion experiments. No fibril-membrane association was detected for either membrane type (**Fig. S6B, C**). Collectively, these findings rule out differences in the intrinsic membrane affinity for α-syn fibrils, while indicating that α-syn monomers bind more strongly to Neuron membranes across all L/P ratios. The distinct fibril-membrane associations observed are therefore likely to originate from protein-membrane interactions occurring during the early stages of membrane-assisted aggregation.

### Conformation-Dependent Intraneuronal Aggregation Properties and Inflammatory Responses

Preformed fibrils (PFFs) were generated by sonicating mature fibrils to examine conformation-dependent pathological effects. PFFs generated from α-syn fibrils grown in the absence of membranes or in the presence of Neuron or Aged membranes are called PFFs, N-PFFs, Aged-PFFs, respectively. To evaluate conformation-dependent prion-like behavior, 0.4 mol% (monomer equivalent) of PFFs, N-PFFs, or Aged-PFFs was added to a 200 μM solution of monomeric α-syn in the absence of membranes, and fibril formation was monitored using ThT fluorescence. Aggregation kinetics were quantified using t_0.1_ and t_0.5_. All PFFs significantly accelerated fibril formation, with N-PFFs and Aged-PFFs showing a trend toward faster fibril formation compared to lipid-free PFFs (**Fig. S11**).

Next, we examined how these polymorphic α-syn fibrils shape disease-relevant phenotypes in dopaminergic neurons, the cell type most affected in PD^82–84^. To this end, we used an in vitro cellular model of midbrain tissue by differentiating neuroblastoma cells, which are commonly employed as a dopaminergic cell model in PD research^85–88^. Specifically, neuroblastoma cells were introduced into 48-well plates and cultured until reaching confluency (**Fig. S12A**). Neuronal differentiation was then induced through sequential application of two distinct differentiation media formulations designed to drive morphological maturation, as previously described^89,90^ (**Fig. S12B**). During the early phase of differentiation (Day 2), cells began to extend thin projections (**Fig. S12C, Day 4**), which progressively developed into a dense network of neurite-like structures by Day 12 (**Fig. S12C, Day 12**). Following differentiation, the cells were evaluated for neuronal features by immunostaining for dendritic and axonal markers. This analysis revealed robust expression of microtubule-associated protein 2 (MAP2) and growth-associated protein 43 (GAP43), which serve as markers for dendrites and axons, respectively, confirming that the differentiated neuroblastoma cells had acquired characteristic neuronal phenotypes^91–95^ (**Fig. S12D**). Their dopaminergic identity was also validated by immunofluorescent staining for tyrosine hydroxylase (TH), an enzyme essential for dopamine biosynthesis and a well-established marker of midbrain dopaminergic neurons^96,97^ (**Fig. S12D)**.

Building upon this neuronal platform, we next investigated how dopaminergic neurons respond to conformationally distinct α-syn PFFs. Specifically, our dopaminergic neuronal cells were treated with PFFs, N-PFFs, and Aged-PFFs to induce pathological features characteristic of PD. To distinguish conformation-dependent effects from those arising from associated lipid components present in N-PFFs and Aged-PFFs, lipid-free PFFs were administered either alone or together with Neuron (PFFs + N-lipid) or Aged membranes (PFFs + Aged-lipid). In addition, α-syn monomers combined with Neuron- or Aged membranes (Mono + N-lipid and Mono + Aged-lipid) were included as controls, while untreated cultures served as the negative control. For all treatments, α-syn species were introduced at a concentration of 1.0 μM (monomer equivalent), consistent with prior studies showing that concentrations between 0.25 and 2.0 μM effectively induce PD-associated pathologies in neuronal cultures without causing acute toxicity^98–104^. Accordingly, all lipid-containing samples (N-PFFs, Aged-PFFs, PFFs + N-lipid, PFFs + Aged-lipid, Mono + N-lipid, and Mono + Aged-lipid) contained 10 μM of membranes, corresponding to an L/P ratio of 10.

After 3 days of treatment, we investigated whether the polymorphic α-syn PFFs differed in their prion-like capacity to template de novo formation of intracellular α-syn aggregates, a pathological hallmark of PD. As phosphorylation of α-syn at Ser129 (pS129) is a defining molecular signature of the pathogenic α-syn assemblies in synucleinopathies^105–107^, we first tested whether distinct PFF conformers differentially promoted pS129 accumulation.

Consistent with their prion-like activity, all PFF treatment groups produced a significant increase in pS129 immunofluorescence compared with untreated controls (**Fig. 6A, B**). Although these differences did not reach statistical significance, conformationally distinct PFFs showed a trend toward differential levels of pS129 accumulation (**Fig. 6B**). Notably, Neuron- and Aged membranes alone did not alter pS129 expression, as indicated by the absence of significant changes in Mono + N-lipid and Mono + Aged-lipid groups relative to untreated controls (**Fig. 6B**). This result was further supported by the observation that addition of Neuron- or Aged membranes to lipid-free PFFs did not affect pS129 levels, as PFFs + N-lipid and PFFs + Aged-lipid exhibited pS129 accumulation similar to lipid-free PFFs alone (**Fig. 6B**).

**Figure 6.**
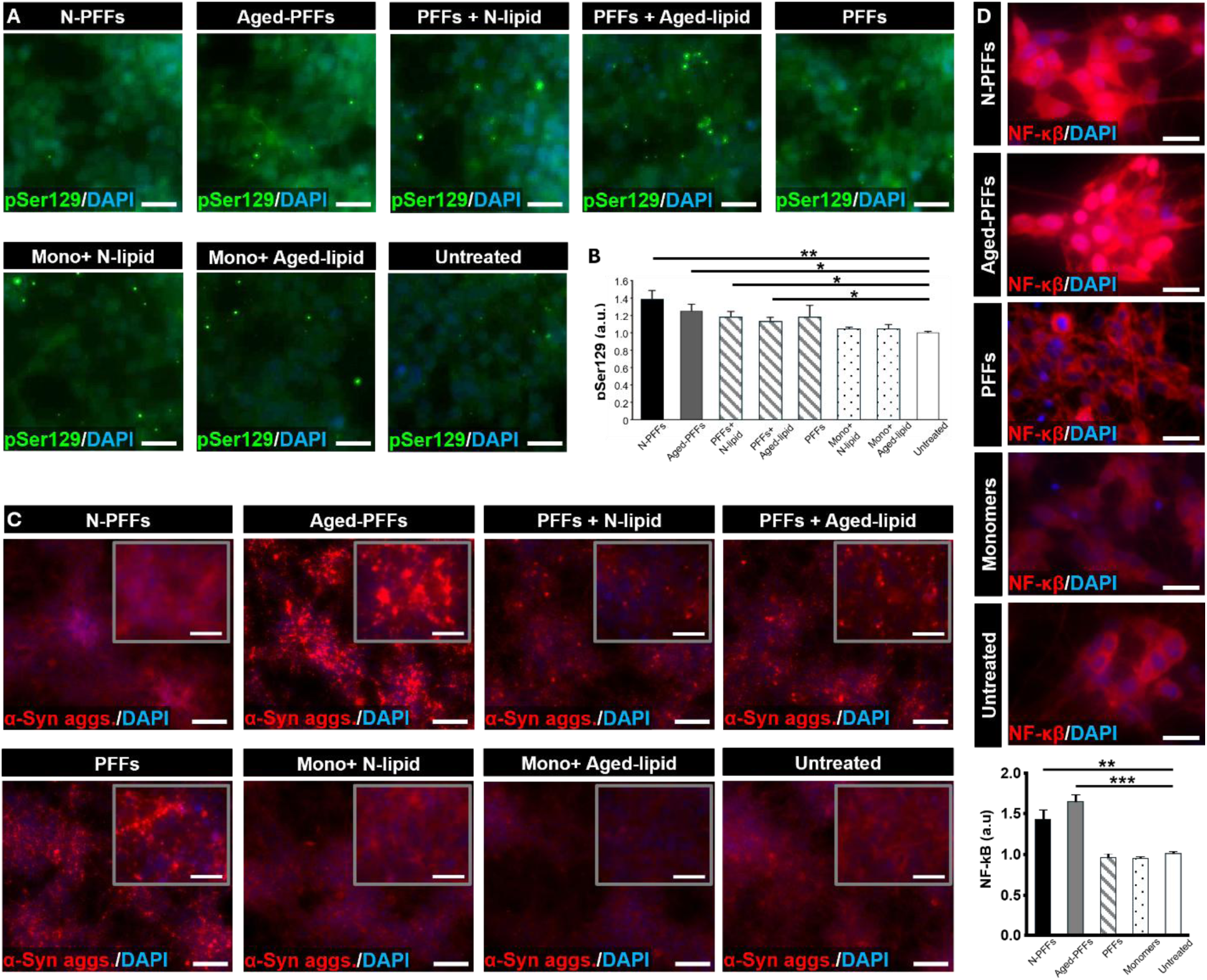
In vitro study on the pathological effects of polymorphic α-syn PFFs in dopaminergic neuronal cells. (A) Immunostaining of phosphorylated α-syn (pS129) tended to show varying levels of endogenous α-syn misfolding following treatments with conformationally distinct α-syn PFFs. Neuron- and Aged-membranes didn’t significantly affect pS129 accumulation. Scale bars, 20 µm. (B) Quantification of pS129 levels in response to distinct α-syn PFF conformations with or without membranes. (C) Immunofluorescence micrographs showing differential formation of intraneuronal α-syn aggregates in response to structurally distinct α-syn PFFs and lipid conditions. Insets show magnified views of α-syn puncta within dopaminergic neurons. Scale bars, 100 µm for main images and 25 µm for insets. (D) Cells treated with distinct α-syn PFFs exhibit differential activation and nuclear translocation of NF-κB. Scale bars, 20 µm. Data show mean ± SD with n = 3. ***P < 0.001, **P < 0.01, *P < 0.05.

Considering that pS129 has been reported to promote α-syn aggregation^105,107,108^, we next examined whether distinct α-syn PFF conformers differentially induced intracellular α-syn aggregate formation in our dopaminergic neuronal cells. To assess this, we immunofluorescently labeled intracellular α-syn fibrils as a readout of punctate α-syn structures, which represent small, discrete aggregates of misfolded α-syn protein. Aged-PFFs and lipid-free PFFs produced abundant intraneuronal α-syn puncta in dopaminergic neuronal cells, with Aged-PFFs yielding larger and more intense punctate aggregates (**Fig. 6C**). This qualitative difference suggests conformational variation among α-syn fibrils that modulates intracellular templating and aggregate expansion (**Fig. 6C**). In contrast, N-PFF-treated cells exhibited little punctate α-syn staining (**Fig. 6C**). Notably, PFFs + Aged-lipid treatment exhibited puncta similar to those observed with lipid-free PFFs, while addition of Neuron membranes to PFFs (PFFs + N-lipid) resulted in qualitatively fewer discernible punctate structures (**Fig. 6C**). Although further analysis is required, these observations suggest that Neuron membranes may suppress punctate α-syn aggregation despite their strong capacity to promote Ser129 phosphorylation. Importantly, control groups, with or without lipid components, did not exhibit detectable α-syn puncta, indicating that lipid components alone are insufficient to induce α-syn aggregation.

In the subsequent phase of the study, we evaluated whether distinct conformations of α-syn PFFs differentially triggered inflammatory responses in neuronal cells, given increasing evidence that α-syn aggregates contribute to neuroinflammation associated with PD progression^109–113^. As a representative readout of neuroinflammatory activation, we analyzed the subcellular localization of nuclear factor-kappa B (NF-κB), a key transcription factor involved in inflammation and cellular stress responses in neurodegeneration^113,114^. As the previous aggregation analyses with lipid controls established that lipid components alone do not drive pathological α-syn aggregation, we simplified the experimental design and focused subsequent inflammation analyses on biologically relevant fibril conformers and baseline controls including Aged-PFFs, N-PFFs, lipid-free PFFs, α-syn monomers, and untreated controls. In the absence of pathogenic PFFs, as in untreated and monomer-treated cultures, NF-κB was largely retained in the cytoplasm (**Fig. 6D**), consistent with its inactive state under non-inflammatory conditions. Exposure to N-PFFs or Aged-PFFs, however, induced robust NF-κB nuclear translocation, as evidenced by increased nuclear and decreased cytoplasmic staining (**Fig. 6D**). Quantitative analysis revealed approximately a 1.5-fold increase in NF-κB nuclear translocation in cultures treated with N-PFFs and Aged-PFFs compared to untreated controls and α-syn monomer-treated cells (**Fig. 6D**). It should be noted that treatment with PFFs did not alter NF-κB distribution, indicating the conformational specificity of the inflammatory response.

Collectively, these in vitro study results show that Aged-PFFs reproduce several hallmark features of PD, including abnormal α-syn phosphorylation, intracellular fibril aggregation, and NF-κB-mediated neuroinflammation, whereas N-PFFs and lipid-free PFFs induce only partial responses and α-syn monomers do not elicit these effects. This suggests that structural variation among α-syn fibril conformers may play a critical role in modulating neuroinflammatory responses as well as protein misfolding and aggregation, key contributors to dopaminergic neurodegeneration in PD.

## Discussion

α-syn membrane binding has been recognized as a critical step in its pathological role, primarily by nucleating aggregation^36,115^. To date, extensive studies have examined how lipids—particularly headgroup composition—affect α-syn aggregation kinetics, largely using simplified membranes composed of one or two lipid types ^25,35–38,40,45^. However, the influence of lipid fatty acid composition, which primarily governs membrane fluidity and packing, on α-syn aggregation within physiologically relevant neuronal membranes remains underexplored. Notably, aging is associated with a decrease in PUFAs and an increase in monounsaturated fatty acids, representing major alterations in brain lipid composition that reduce membrane fluidity and dynamics^50–52^. Here, we provide important insights into how these age-related alterations in membranes influence α-syn fibrils and their pathological effects by using two neuronal membrane models that recapitulate normal and aged lipid compositions.

Our ssNMR results demonstrate that α-syn adopts structurally distinct fibrils when it aggregates in the presence of membranes with different lipid compositions, and the resulting fibrils exhibit distinct membrane-association properties. These membrane-association differences likely arise from variations in membrane-monomer interactions (**Fig. S6A**). Specifically, α-syn fibrils formed with Aged membranes exhibit weaker membrane association compared to those formed with Neuron membranes. To verify that this weaker interaction was not due to the relatively low lipid content at an L/P ratio of 10, we performed 2D ^13^C-detected ^1^H-^1^H spin diffusion experiments on α-syn fibrils grown with Aged membranes at an L/P ratio of 50. The lipid-to-protein signal intensity was comparable to that observed at an L/P ratio of 10 (**Fig. S7**), confirming that the reduced lipid-to-protein signals are not attributable to the lower amount of lipids.

Membranes composed of zwitterionic lipids lacking charged species are known to modulate α-syn binding affinity through lipid packing defects, transient surface gaps between lipid headgroups that expose hydrophobic acyl chains to solvent^116^. The extent of lipid packing defects, which increases with the degree of fatty acid unsaturation, promotes hydrophobic interactions that drive α-syn insertion into the membrane bilayer^116,117^. Thus, the enhanced membrane binding of α-syn monomers to Neuron membranes, which contain more unsaturated lipids (POPC and DOPE) compared to Aged membranes (DPPC and POPE), can be attributed to the higher level of lipid packing defects in Neuron membranes. These distinct monomer-membrane interactions may give rise to different fibril structures. Although thorough comparisons will require the development of structural models for each fibril, ssNMR data suggest that the fibril cores span residues 36-91 for α-syn fibrils grown with Neuron membranes and residues 14-100 for those grown with Aged membranes (**Fig. S5**). This indicates that fibrils formed with Neuron membranes retain a longer N-terminal disordered region (residues from 1-35) than those formed with Aged membranes (residues from 1-13). Because the N-terminal region is known to mediate membrane binding^66,118–120^, its stronger affinity for Neuron membranes likely promotes insertion of a longer portion of this region into the lipid bilayer. This insertion may limit its involvement in fibril core formation through protein-protein hydrophobic interactions, resulting in a longer disordered segment compared to fibrils grown with Aged membranes. However, further studies are needed to determine the molecular structures of these fibrils and to investigate residue-specific membrane binding throughout fibril formation, including monomers, oligomers, and fibrils, to gain deeper insight into the structural polymorphisms of membrane-associated fibrils.

To translate membrane-associated α-syn conformers into a patho-physiologically relevant context, we employed an in vitro cell-based platform to investigate the pathological consequences of α-syn polymorphism. Using this interdisciplinary approach, we found that distinct α-syn PFF conformations elicited differential responses in dopaminergic neuronal cells across multiple pathological processes implicated in PD neurodegeneration. Although all α-syn PFF conformations significantly increased Ser129 phosphorylation relative to controls (**Fig. 6A, B**), their capacities to drive intracellular aggregation differed markedly (**Fig. 6C**). Specifically, N-PFFs induced robust pS129 accumulation but failed to generate discernible α-syn aggregates, whereas Aged-PFFs and lipid-free PFFs produced prominent intraneuronal aggregates. Notably, Aged-PFFs elicited larger and more abundant α-syn aggregates than N-PFFs and lipid-free PFFs (**Fig. 6C**). Both N-PFFs and Aged-PFFs retain Neuron and Aged membranes, respectively, raising the possibility that lipid mediated effects could contribute to the observed cellular responses in addition to fibril conformation. To disentangle lipid-driven effects from fibril-driven effects, we therefore incorporated control conditions in which Neuron or Aged membranes were applied to α-syn monomers and to lipid-free PFFs. Using this expanded set of treatments, we examined α-syn phosphorylation and intraneuronal α-syn aggregation (**Fig. 6A-C**). These experiments showed that lipid components alone do not induce pathological α-syn modification or punctate aggregation. Moreover, Neuron membranes, whether inherently present in N-PFFs or added to lipid-free PFFs (PFFs + N-lipid), do not reproduce the strong aggregate accumulation observed with Aged-PFFs or PFFs + Aged-lipid. Although further validation is needed, these results suggest that Neuron-related lipid components suppress the formation of punctate α-syn aggregates despite strongly promoting Ser129 phosphorylation. This observation may be partly attributed to the higher binding affinity of Neuron membranes for α-syn monomers (**Fig.S2A**), thereby reducing the pool of free monomers available for fibril growth.

Notably, Aged-PFFs induced greater NF-κB nuclear translocation than N-PFFs and lipid-free PFFs (**Fig. 6C, D**). This enhanced inflammatory activation, together with the abundant intraneuronal aggregates induced by Aged-PFFs may provide mechanistic insight into clinical differences between young- and late-onset PD progression. Later-onset PD is typically associated with more severe and rapidly progressing symptoms, including motor deficits, cognitive impairment, and greater dopaminergic dysfunction^121–123^. Given the central role of neuroinflammation in initiating and accelerating neurodegenerative processes^124,125^, the distinct α-syn fibril conformations generated in aged membrane environments may promote more severe and rapid PD progression through enhanced inflammatory responses. However, further studies will be required to determine whether Aged-PFFs specifically drive pathological features characteristic of late-onset PD.

Collectively, these results demonstrate that distinct α-syn PFF conformations differ in their ability to engage multiple pathological pathways. These findings support the idea that membrane-mediated structural diversity of α-syn fibrils contributes to phenotypic heterogeneity in synucleinopathies^10,126,127^. Among them, Aged-PFFs emerged as the most pathologically relevant species, as they uniquely triggered the full spectrum of α-syn pathology examined here, including abnormal phosphorylation, intracellular aggregation, and inflammatory activation. This enhanced disease-driving capacity is consistent with their formation under pathologically relevant conditions, in which α-syn monomers interact with aged neuron membrane-related lipid components. Such lipid environments may more closely emulate the subcellular milieu of aging neurons that modulates α-syn aggregation dynamics and fibril conformation in PD, thereby conferring increased pathological potency.

Although, our membrane models capture the major physiochemical features of normal and aged neuronal membranes, they do not fully recapitulate the compositional complexity of biological membranes. In particular, our models exclude lipid species that individually account for less than ∼10% of total phospholipids, including PUFAs^54,128,129^, due to the simplified four-lipid formulation. Because the most significant age-related changes involve alterations in fatty acid saturation levels largely driven by PUFAs, future studies incorporating these lipid species will be necessary to fully define their contributions to α-syn aggregation and membrane interactions.

Similarly, while our in vitro dopaminergic neuronal model recapitulates several hallmark features of PD pathology, it remains an incomplete representation of the functional complexity of in vivo dopaminergic neurons. In particular, the absence of validated synaptic connectivity may limit its ability to capture interneuronal transmission of α-syn species, a process thought to play a pivotal role in disease progression^130–132^. This limitation constrains the extent to which the cellular model can be used to investigate disease mechanisms, as it does not account for the interneuronal spread of pathogenic α-syn, an important aspect of disease progression. To improve the translational relevance of this cellular model, future efforts will need to demonstrate reliable formation of synaptic structures and the occurrence of action potential-mediated synaptic transmission.

## Conclusion

In conclusion, our study demonstrates that complex membranes, containing cholesterol, phospholipids, and sphingolipids, which mimic neuronal plasma membranes, accelerate α-syn aggregation and generate fibril structures distinct from those formed without membranes. ssNMR data further reveal that age-associated membrane changes in phospholipid fatty acid chains alter both the structures and aggregation kinetics of α-syn fibrils. Moreover, the differences in intraneuronal aggregation and inflammatory responses induced by distinct forms of PFFs, highlight the critical role of membrane composition in shaping pathogenic protein aggregation. Together, these findings advance our understanding of membrane-mediated, conformation-dependent pathologies and provide insights into the interplay between neurodegenerative aggregates and membranes.

## ASSOCIATED CONTENT

### Supporting Information

TEM images showing diverse morphologies of α-syn fibrils, Proteinase K digestion analysis, Full 2D ssNMR spectra of α-syn fibrils, Analysis of distinct α-syn fibril cores, Membrane binding analysis of α-syn monomers and fibrils, Membrane association of α-syn fibrils at L/P = 50, Residue-specific membrane interactions of α-syn fibrils, Distribution of PFF lengths, Calcein membrane leakage assay, Conformation-dependent prion-like activities, Differentiation of neuroblastoma cells into dopaminergic neurons, and Long-term stability of vesicles (PDF).

## AUTHOR INFORMATION

### Corresponding Author

**Myungwoon Lee** – Department of Chemistry, Drexel University, Pennsylvania 19104, United States;

### Author Contributions

M.L and J.P designed the project. Y.B, A.A, Y.D and T.O performed experiments. M.L, J.P, Y.B, and A.A wrote the manuscript.

### Notes

The authors declare no competing financial interest.

## ACKNOWLEDGEMENTS

We thank Drs. Ivan Hung and Zhehong Gan at the National High Magnetic Field Laboratory for their assistance with ssNMR experiments. The National High Magnetic Field Laboratory is supported by the National Science Foundation through NSF/DMR-2128556 and the state of Florida.

We thank Dr. Yury Gogotsi and Alhourani Jamal from the Department of Materials Science and Engineering at Drexel University for their assistance with DLS measurements.

This work was supported by NIH grant NS139178 to M.L and J.P.

