## Supplemental Figures for "Lipid Acyl Chain-Driven α-Synuclein Fibril Polymorphisms and Neuronal Pathologies"

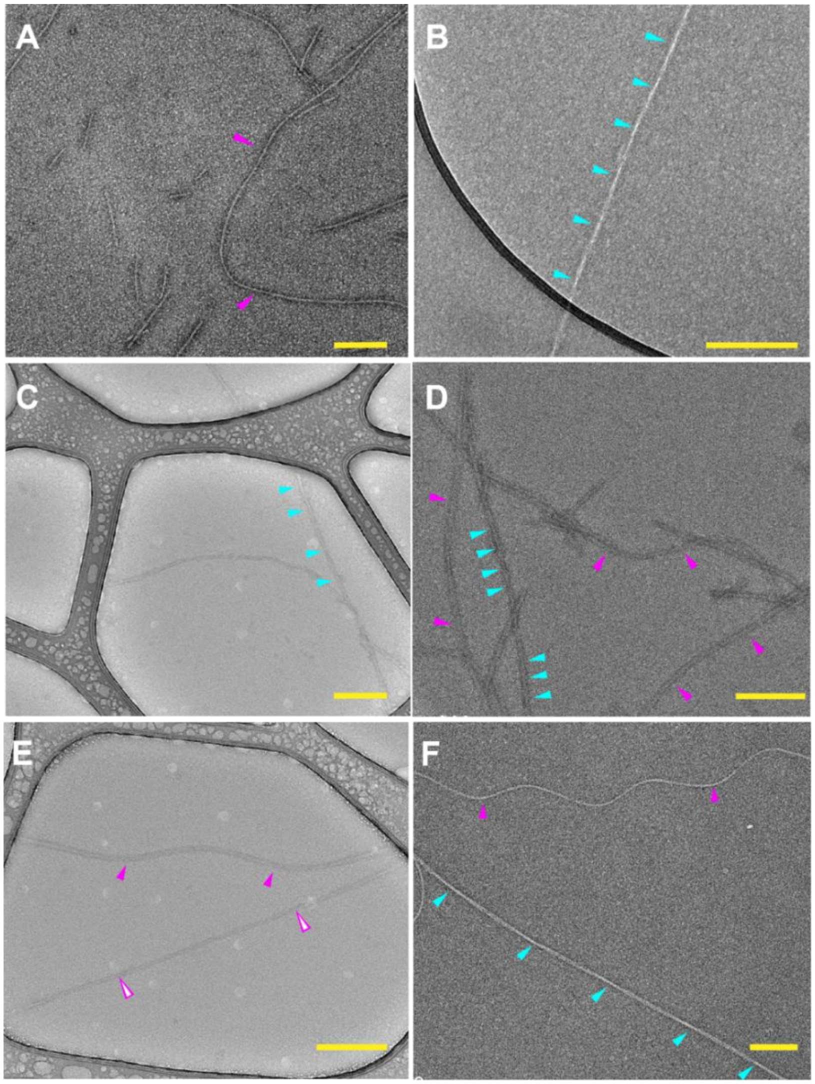


**Figure S1.** Conformational polymorphism of α-syn fibrils within and between samples. (A, B) Lipid-free α-syn fibrils. (C, D) α-syn fibrils formed with Neuron membranes at a L/P ratio of 10. (E, F) α-syn fibrils formed with Aged membranes at a L/P ratio of 10. Twisted and untwisted fibril morphologies are indicated by cyan and magenta arrows, respectively. Open magenta arrows denote thin untwisted fibrils, whereas filled magenta arrows indicate wider untwisted fibrils. Scale bar: 200 nm.

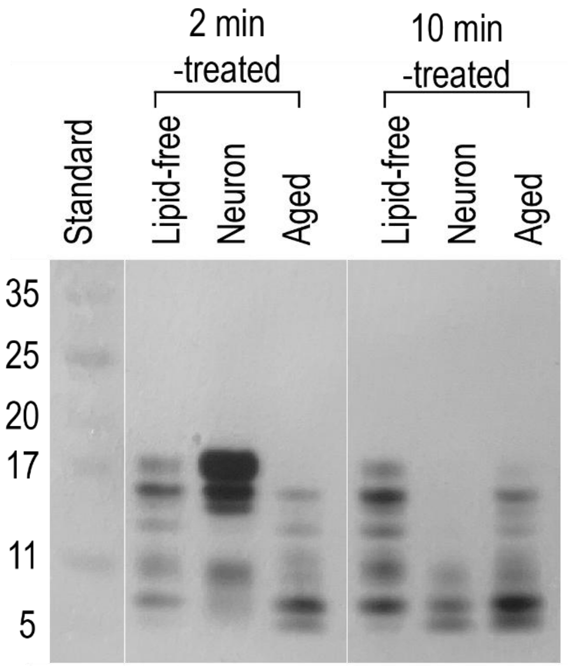


**Figure S2.** SDS-PAGE gel showing α-syn fibrils formed in the absence and presence of membranes after 2- and 10 min proteinase K (PK) digestion.

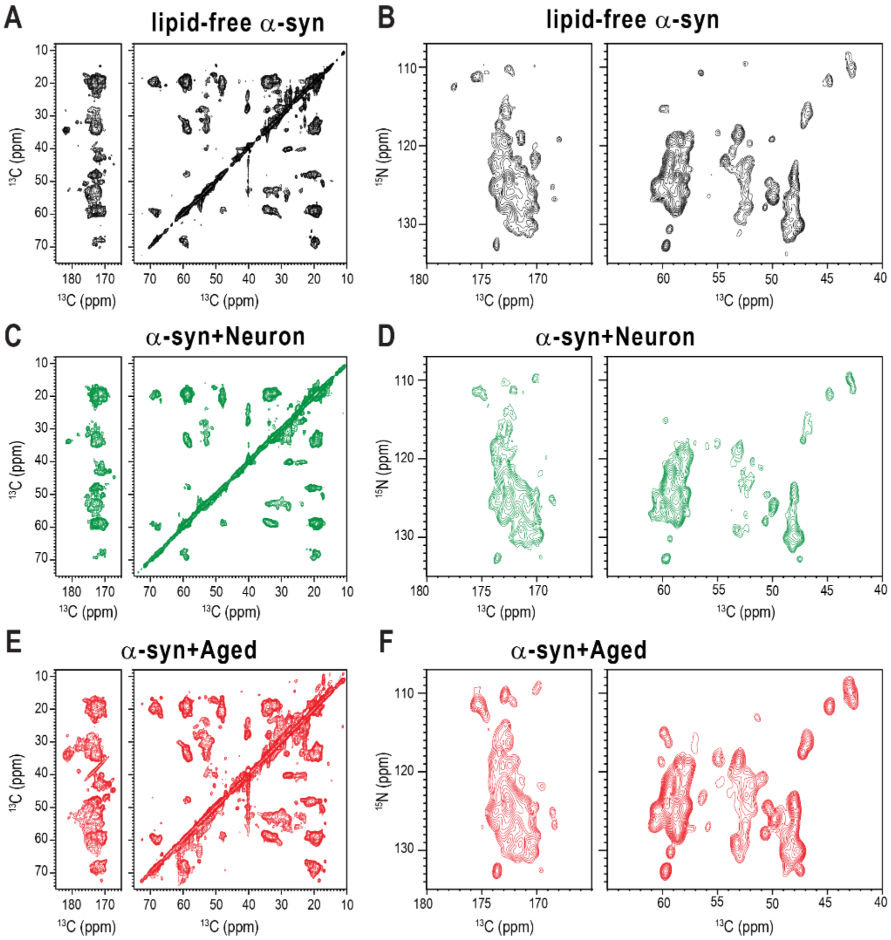


**Figure S3.** Effects of membrane-bound α-syn on fibril structures. 2D $^{13}$C-$^{13}$C (A) and $^{15}$N-$^{13}$C (B) correlation spectra of uniformly $^{13}$C, $^{15}$N-labeled lipid-free α-syn fibrils with a 100 ms DARR mixing time. 2D $^{13}$C-$^{13}$C (C) and $^{15}$N-$^{13}$C (D) correlation spectra of α-syn fibrils grown in the presence of Neuron membrane with a 50 ms DARR mixing time. 2D $^{13}$C-$^{13}$C (E) and $^{15}$N-$^{13}$C (F) correlation spectra of α-syn fibrils grown in the presence of Aged membrane with a 50 ms DARR mixing time.

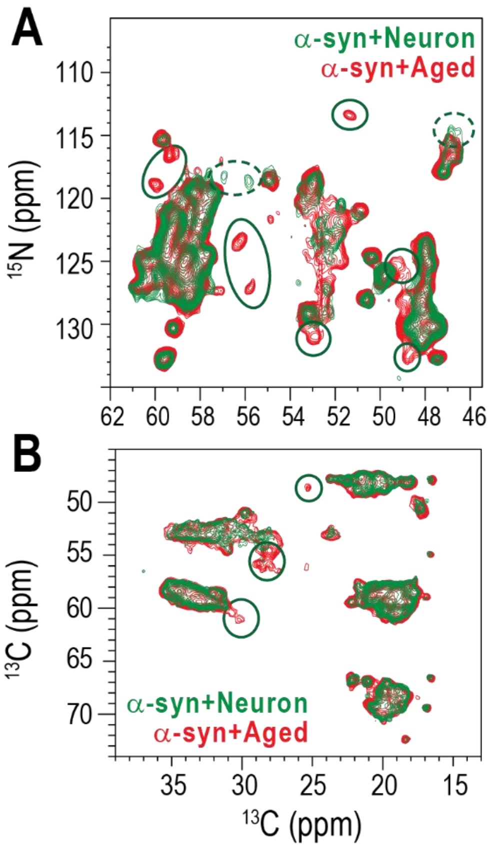


**Figure S4.** Comparison of 2D NMR spectra of α-syn fibrils formed under different membrane conditions. 2D $^{15}$N-$^{13}$C (A) and 2D $^{13}$C-$^{13}$C (B) correlation spectra of α-syn fibrils formed with Neuron membranes (green) overlaid with those formed with Aged membranes (red). Dotted green circles indicate peaks present only in α-syn fibrils grown with Neuron membranes, whereas solid green circles highlight peaks unique to α-syn fibrils grown with Aged membranes.

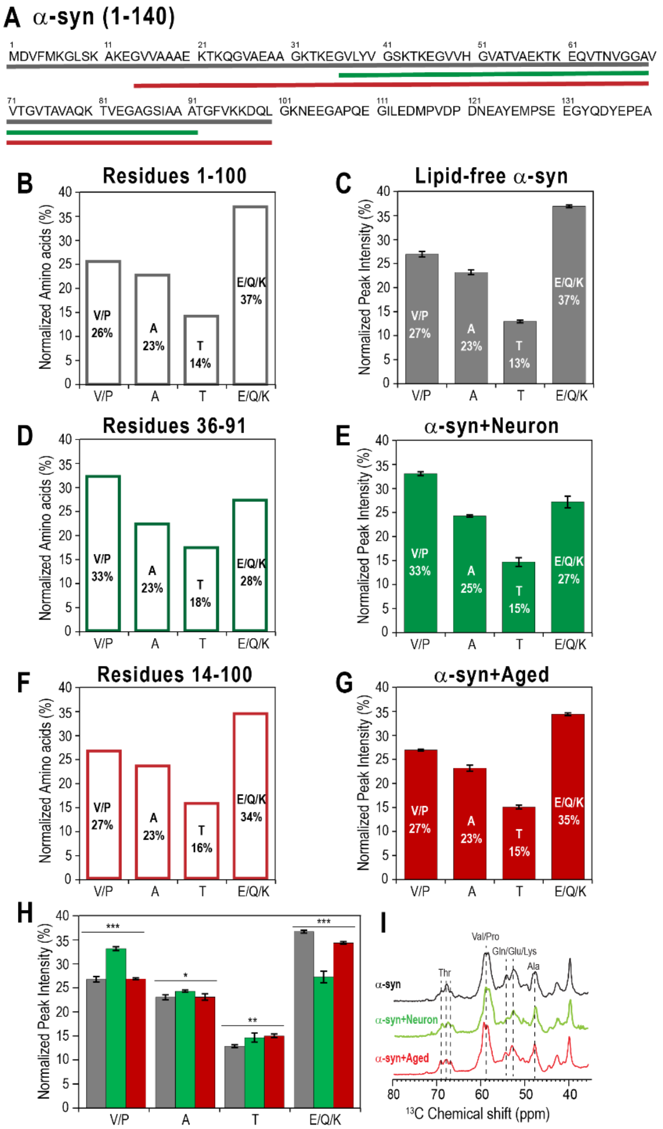


**Figure S5**. Distinct α-syn fibril cores formed under lipid-free conditions and in the presence of Neuron or Aged membranes. (A) Sequence of the α-syn protein (residues 1-140), with color-coded regions indicating expected fibril core variations. Possible fibril cores span residues 1-100 (gray) in lipid-free fibrils, 38-95 (green) in fibrils formed with Neuron membranes, and 14-100 (red) in fibrils formed with Aged membranes. (B) Normalized amino acid count within residues 1-100. (C) Normalized peak intensities of amino acids from the 2D $^{13}$C-$^{13}$C spectrum of lipid-free α-syn fibrils. (D) Normalized amino acid count within residues 38-95. (E) Normalized peak intensities of amino acids from the 2D $^{13}$C-$^{13}$C spectrum of α-syn fibrils grown with Neuron membranes. (F) Normalized amino acid count within residues 14-100. (G) Normalized peak intensities of amino acids from the 2D $^{13}$C-$^{13}$C spectrum of α-syn fibrils grown with Aged membranes. (H) Quantification of residue-specific Cα-Cβ cross peak intensities from the 2D $^{13}$C-$^{13}$C spectra, comparing the relative amino acid composition of fibril core among lipid-free α-syn (gray), α-syn + Neuron (green), and α-syn + Aged (red) fibrils. (I) 1D $^{13}$C CP spectra of lipid-free α-syn fibrils, α-syn fibrils formed with Neuron, and with Aged membranes. Data show mean ± SD with n = 3. ***$P < 0.001$, **$P < 0.01$, *$P < 0.05$.

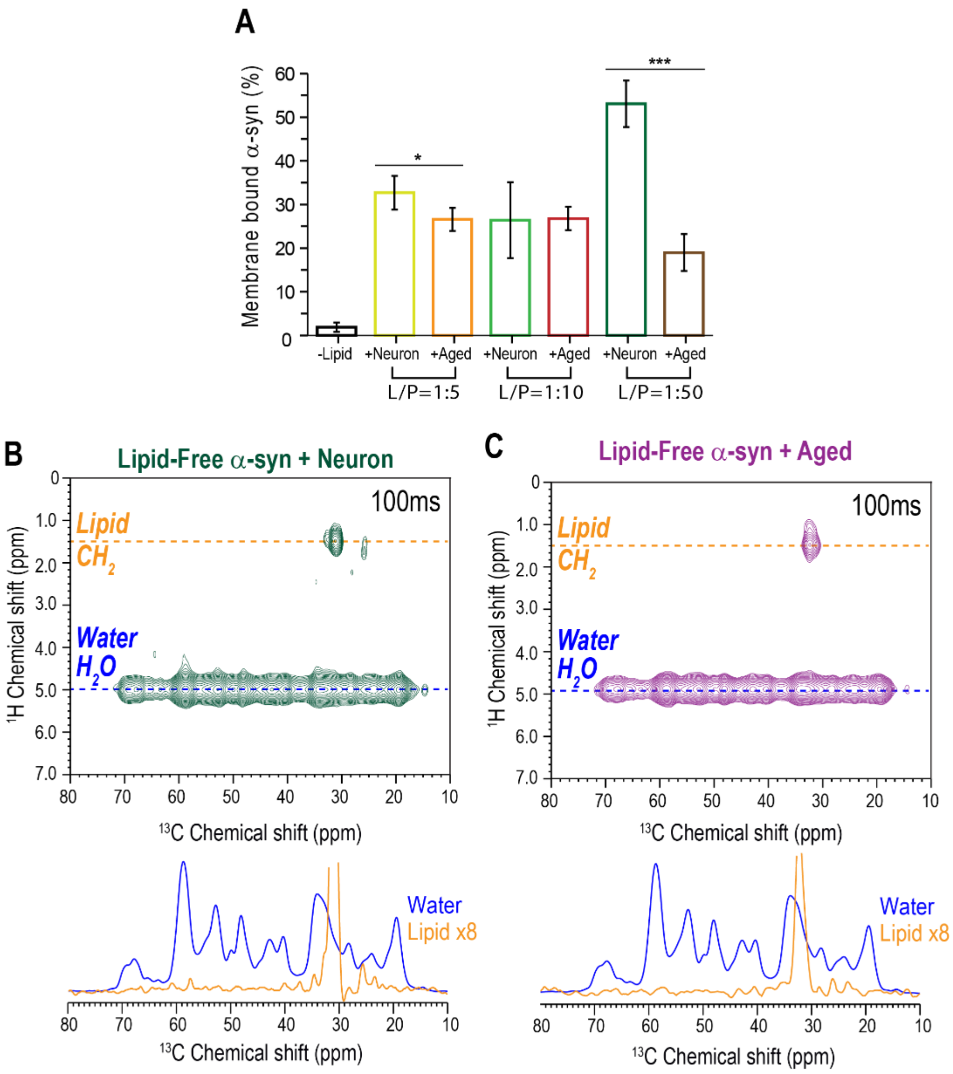


**Figure S6**. (**A**) Membrane binding of α-syn monomer to Neuron and Aged lipid vesicles. Percentages of aggregated or membrane-bound α-syn are shown as bar graphs. Membrane-bound α-syn was measured in the presence of Neuron membranes at L/P ratios of 5 (lime green), 10 (green), and 50 (dark green), and Aged membranes at L/P ratios of 5 (orange), 10 (red), and 50 (brown). Data show mean ± SD with n = 4. ***P < 0.001, **P < 0.01, *P < 0.05. (B, C) Representative 2D $^{13}$C-detected $^{1}$H-$^{1}$H spin diffusion NMR spectra of lipid-free α-syn fibrils mixed with Neuron membranes (B) and with Aged membranes (C) measured at 100 ms. $^{13}$C cross-sections at $^{1}$H chemical shifts of lipid acyl chain (1.3 ppm, orange) and water (5.0 ppm, blue), extracted from 2D $^{1}$H-$^{13}$C spectra measured at $^{1}$H-$^{1}$H spin diffusion mixing times of 4 ms and 100 ms. No protein signals transferred from lipids in either lipid-free α-syn fibrils mixed with Neuron or Aged membranes.

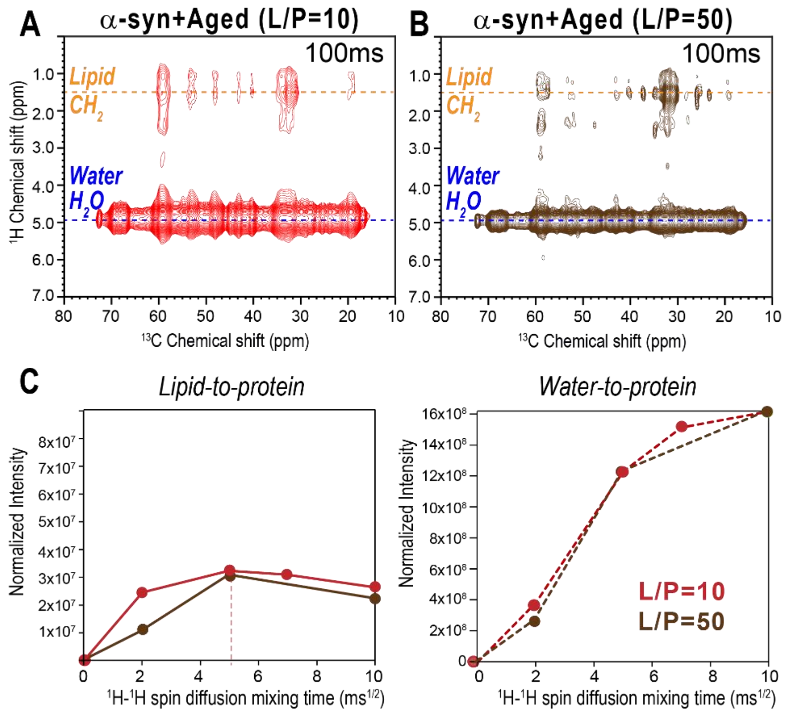


**Figure S7.** Membrane association of α-syn fibrils at low (L/P = 10) and high (L/P = 50) membrane ratios. (A, B) Representative 2D $^1$H-$^{13}$C correlation spectra of α-syn fibrils grown with Aged membranes at L/P = 10 (A) and L/P = 50 (B), measured with a $^1$H-$^1$H spin diffusion mixing time of 100 ms. (C) Lipid-to-protein (solid lines, left) and water-to-protein (dotted lines, right) $^1$H polarization transfer curves as a function of $^1$H mixing time. For both membrane ratios, lipid-derived protein signals plateau after ~25 ms, indicating no enhanced lipid-protein interactions at higher membrane concentrations.

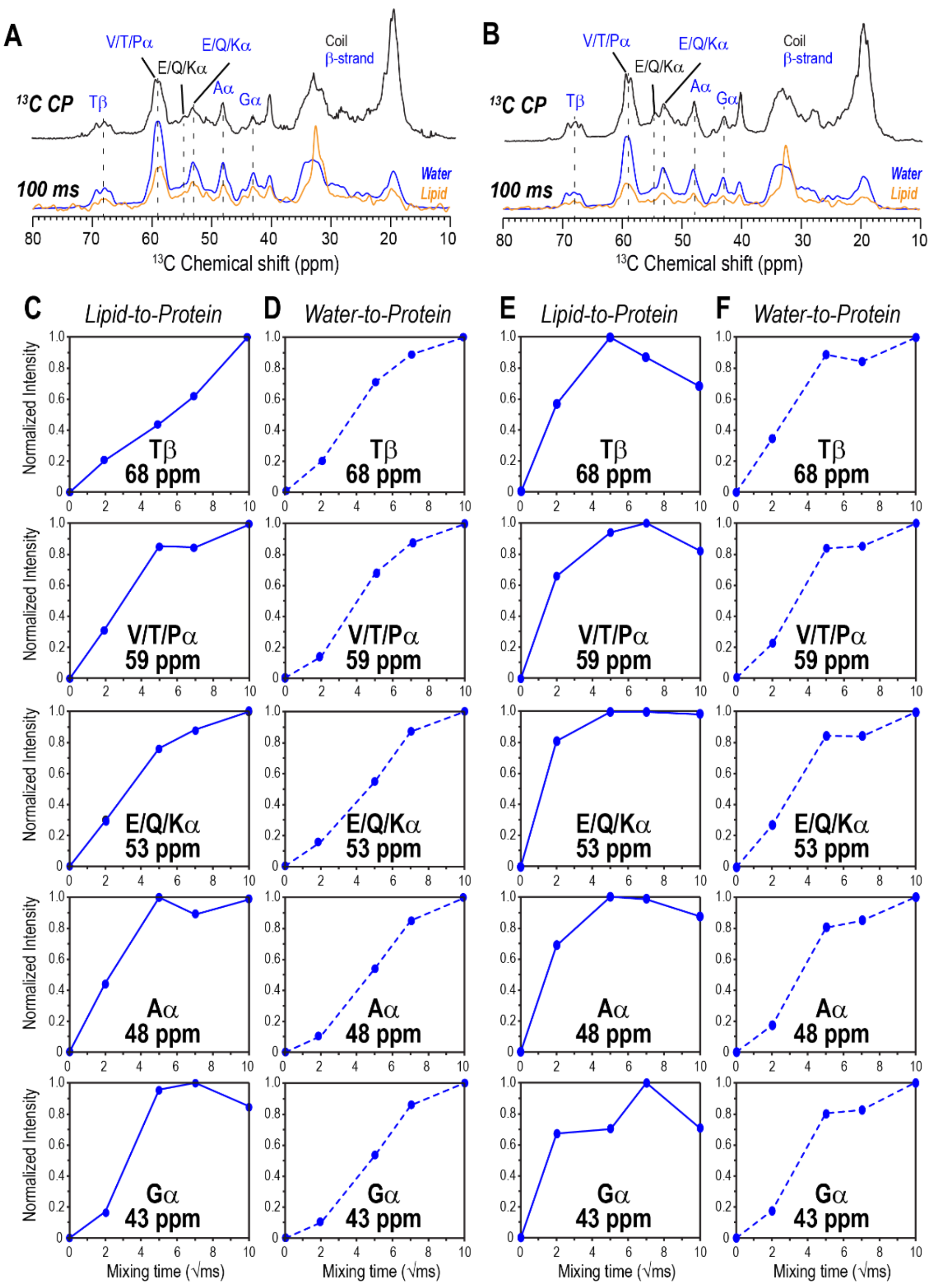


**Figure S8.** Interactions of α-syn fibrils with Neuron or Aged membranes. (A, B) 1D $^{13}$C CP spectra and $^{13}$C cross-sections at water (blue) and lipid $CH_2$ (orange) $^{1}$H chemical shifts, extracted from 2D $^{1}$H-$^{13}$C correlation spectra measured at a 100 ms $^{1}$H mixing time, comparing α-syn fibrils associated with Neuron (A) and Aged (B) membranes. Labeled residues indicate β-sheet (blue) and coil (black) conformations. (C, E) Normalized lipid-to-protein intensity build-up curves for labeled residues in (A) and (B) as a function of $^{1}$H mixing time, extracted from 2D $^{1}$H-$^{13}$C correlation spectra of α-syn fibrils formed with Neuron (C) and Aged (E) membranes. Aged membrane-associated fibrils show signal decay at 100 ms, suggesting weaker lipid-protein interactions. (D, F) Normalized water-to-protein intensity build-up curves for labeled residues in (A) and (B) as a function of $^{1}$H mixing time, extracted from 2D $^{1}$H-$^{13}$C correlation spectra of α-syn fibrils formed with Neuron (D) and Aged (F) membranes.

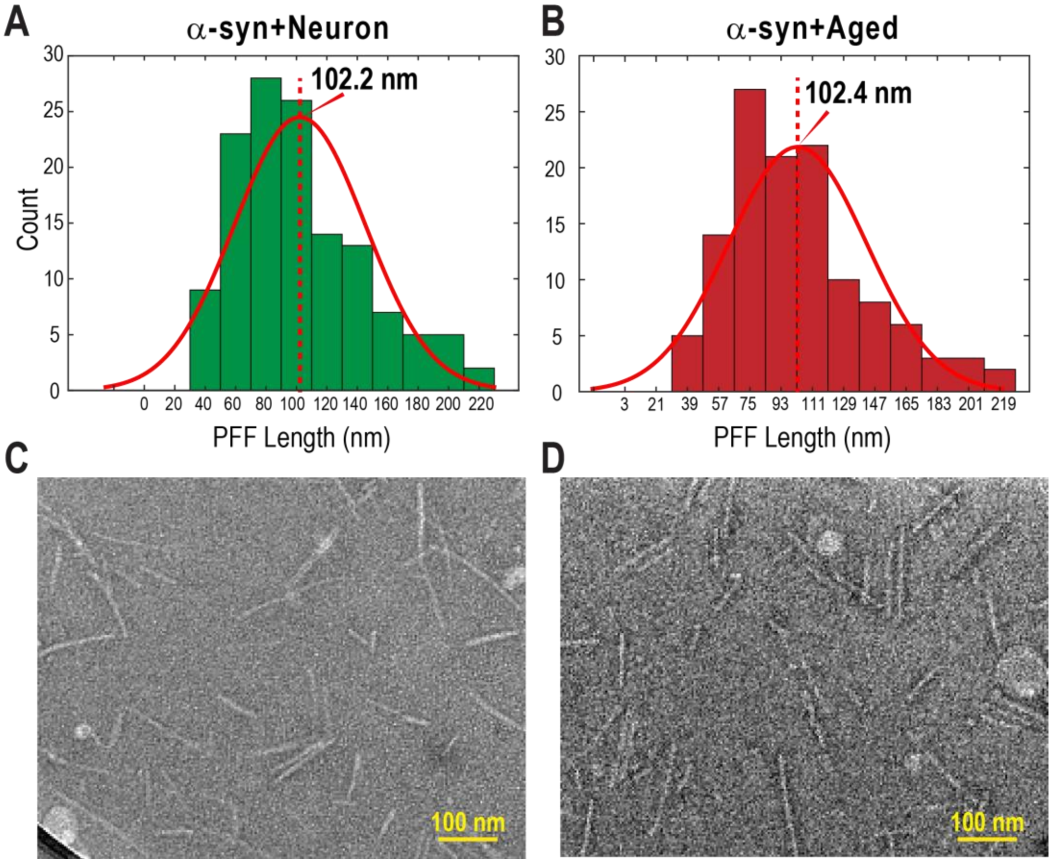


**Figure S9.** Distribution of PFF lengths. (A, B) Histograms of α-syn PFFs grown with Neuron (A) and Aged membranes (B) from negatively stained TEM images. Red lines represent Gaussian fits centered at 102.2 nm and 102.4 nm, respectively. (C, D) Negatively stained TEM images of α-syn PFFs grown with Neuron (C) and Aged (D) membranes showing uniform fibril lengths. Scale bars represent 100 nm.

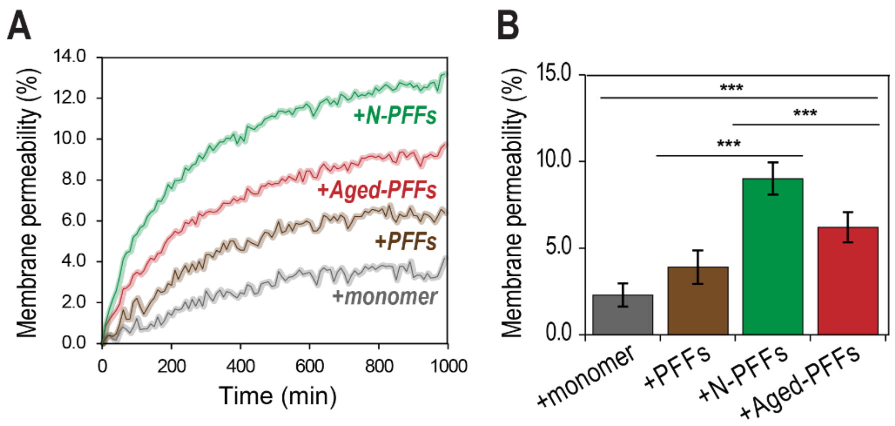


**Figure S10.** Membrane permeability measured using a calcein leakage assay. (A) Time-dependent calcein release from Neuron vesicles exposed to α-syn monomers (gray), lipid-free PFFs (PFFs), or PFFs grown with Neuron (N-PFFs, green) or Aged (Aged-PFFs, red) membranes. (B) Quantification of calcein leakage shown as a bar graph. N-PFFs caused the strongest membrane permeability and disruption, followed by Aged-PFFs-treated vesicles. Data are presented as mean ± SD with n = 3. ***P < 0.001, **P < 0.01, *P < 0.05.

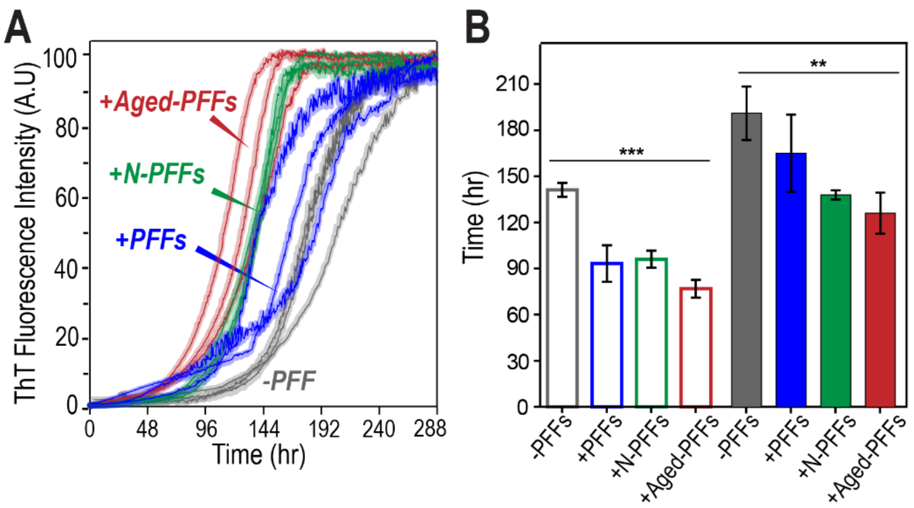


**Figure S11.** Conformation-dependent prion-like seeding activity of lipid-free PFFs, N-PFFs, and Aged-PFFs. (A) ThT fluorescence curves of α-syn aggregation kinetics at monomer concentration of 200 μM in the absence and presence of various PFFs: in the absence of PFFs (gray), in the presence of 0.4 mol% (monomer equivalent) of lipid-free PFFs (blue), N-PFFs (green), and Aged-PFFs (red) over 288 hours. (B) Quantitative comparison of aggregation kinetics using $t_{0.1}$ (open bars) and $t_{0.5}$ (solid bars), representing the times required to reach 10% and 50% of maximal ThT fluorescence intensity, respectively. The same color scheme as in (A) is used. Data are presented as mean ± SD with n = 3. ***$P < 0.001$, **$P < 0.01$, *$P < 0.05$.

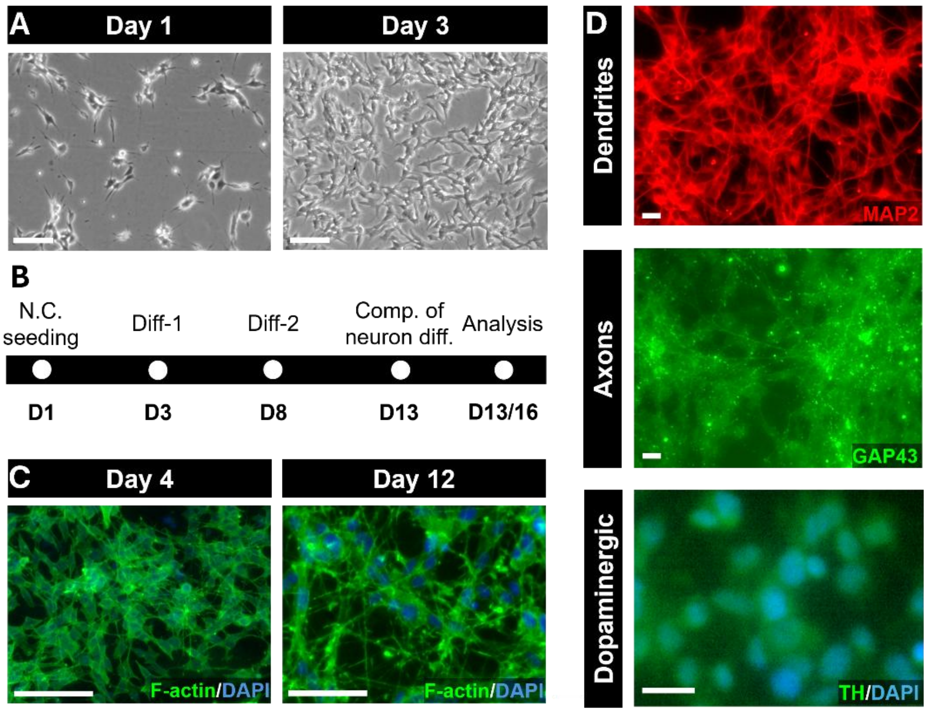


**Figure S12.** Differentiation of neuroblastoma cells into dopaminergic neurons. (A) Neuroblastoma cell seeding and culture confluency after 2 days. Scale bars, 100 µm. (B) Timeline of dopaminergic differentiation in neuroblastoma cells. Abbreviations: N.C. (neuron chamber), D (day), comp.(completion), and diff. (differentiation). (C) Phalloidin staining shows morphological changes in neuroblastoma cells during differentiation at Day 4 and Day 12. Scale bars, 100 µm. (D) Immunofluorescence staining micrographs confirm successful differentiation of neuroblastoma cells into dopaminergic neuronal cells after 13 days. MAP2 and GAP43 staining reveal dendritic and axonal structures, respectively, while tyrosine hydroxylase (TH) staining confirms dopaminergic identity. Scale bars, 20 µm.

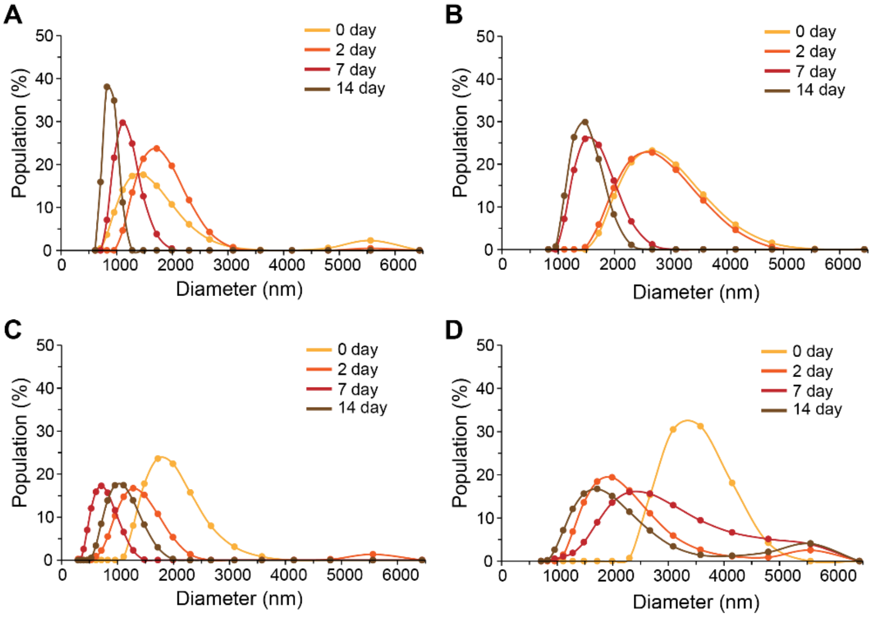


**Figure S13.** Long-term stability of Neuron and Aged membranes in the absence and presence of α-syn monitored by DLS. Vesicle diameter distributions were measured over a 14-day (336-h) incubation at 37 °C for 2 mM Neuron membranes (A), 2 mM Aged membranes (B), Neuron membranes with 200 μM α-syn monomers (L/P = 10) (C), and Aged membranes with 200 μM α-syn monomers (L/P = 10) (D).
